# Histone neutralization protects the ischemic brain against stroke-associated pneumonia

**DOI:** 10.64898/2026.02.28.708679

**Authors:** Dongpei Yin, AnRan Li, Olga Shevchuk, Ayan Mohamud Yusuf, Janine Gronewold, Stephanie Thiebes, Tobias Tertel, Chen Wang, Nina Hagemann, Yiqiao Zhang, Claudia Graser, Humeyra Tas, Michael Fleischer, Britta Kaltwasser, Benedikt Frank, Ali Ata Tuz, Vikramjeet Singh, Devon Siemes, Ekaterina Pylaeva, Bente Siebels, Hartmut Schlueter, Jadwiga Jablonska, Yi Liu, Fengyan Jin, Egor Dzyubenko, Jens Minnerup, Luisa Klotz, Bernd Giebel, Oliver Soehnlein, Matthias Gunzer, Daniel R. Engel, Dirk M. Hermann

## Abstract

Bacterial pneumonia aggravates ischemic stroke via mechanisms that still remain to be determined. In ischemic stroke patients and mice exposed to middle cerebral artery occlusion, we show that stroke-associated pneumonia markedly worsens clinical stroke outcome. In mice, pneumonia induced 3 days after stroke impaired neurological recovery and increased brain neutrophil infiltrates, blood-brain barrier breakdown, cerebral microvascular thrombosis, and progressive brain atrophy. The antibiotic amoxicillin only partially ameliorated pneumonia-associated neurological deficits and neutrophil infiltrates. Neutrophils were critical mediators of pneumonia-induced blood-brain barrier breakdown and microvascular thrombosis. Notably, administration of a neutralizing anti-histone antibody during pneumonia—unlike degradation or blockade of neutrophil extracellular trap formation or myeloperoxidase inhibition—restored long-term neurological recovery and prevented brain atrophy in stroke-associated pneumonia mice. This study identifies extracellular histones as key drivers of secondary inflammatory brain injury and establishes histone neutralization as a therapeutic strategy with an extended treatment window in the post-acute stroke phase.

**One Sentence Summary:** Neutralizing extracellular histones reverses pneumonia-driven secondary brain injury and restores long-term recovery after ischemic stroke.

## INTRODUCTION

Despite recent progress in recanalizing therapies, ischemic stroke remains the leading cause of long-term disability and second leading cause of death worldwide ^1^. In view of this disease burden, there is a major need of treatments that mitigate the long-term consequences of ischemic stroke and enhance recovery in the post-acute stroke phase^2–4^. Neurorestorative treatments so far failed in randomized controlled clinical trials ^5–7^. Ischemic stroke can trigger various complications, including infections and thrombosis ^1,8^. Due to post-ischemic immunosuppression, ∼30% of stroke patients suffer infections in acute hospital settings ^8^. Pneumonia incidence alone is ∼8.5-14.3%, and pneumonia incidence of stroke patients on intensive care units is 28% ^8^. Pneumonia predisposes to poor stroke outcome: Via the development of sepsis, pneumonia is a leading cause of stroke-associated deaths ^9,10^.

Several studies have explored prophylactic antibiotics to prevent stroke-associated pneumonia ^11,12^. In these studies, prophylactic antibiotics consistently did not yield beneficial effects on stroke outcome ^11,12^. Yet, physician-diagnosed pneumonia was independently associated with unfavorable 3-month outcome in one large randomized study ^12^, while pneumonia diagnosed based on a predefined algorithm was not associated with unfavorable outcome in another one ^11^. Some studies have administered β-blockers or interferon-γ to alleviate stroke-induced immunosuppression ^13,14^. Both treatments again did not prevent stroke-associated pneumonia nor enhance stroke outcome ^13,14^. The question whether pneumonia increases stroke damage remains open and underlying mechanisms are still to be explored.

Polymorphonuclear neutrophils are the most abundant leukocytes in humans, which accumulate in the brain parenchyma within hours after ischemic stroke ^15,16^. We previously showed that neutrophils exaggerate ischemic injury and neurological outcome in mice exposed to transient middle cerebral artery occlusion (MCAO) ^15^. Recent studies have found that activated neutrophils can release neutrophil extracellular traps (NETs), web-like structures composed of DNA, to which proteins such as myeloperoxidase (MPO) or histones are bound ^17^. NETs contribute to host defense by capturing bacterial pathogens ^18^. NETs have recently been shown to mediate post-stroke immunosuppression ^19^. The role of neutrophils and their secreted contents for neurological outcome in stroke-associated pneumonia was unknown.

We herein asked whether and how stroke-associated pneumonia influences neurological recovery and ischemic injury. Thus, we prospectively examined consequences of stroke-associated pneumonia for neurological recovery in ischemic stroke patients within the “Neutrophils: Origin, Fate and Function” Stroke (NOFF-S) cohort and evaluated the effects of bacterial pneumonia induced by intratracheal *S. pneumoniae* instillation on neurological recovery and ischemic injury in middle cerebral artery occlusion (MCAO) mice. *S. pneumoniae* is the principal cause of bacterial pneumonia in developed and developing countries that is responsible for more infection-related human deaths than any other bacterium ^20^. *S. pneumoniae* is one of the most common pathogens in stroke-associated pneumonia. It was therefore the pathogen of choice for us for our experimental studies. After showing that bacterial pneumonia exacerbates neurological outcome and ischemic injury in MCAO mice similar to neurological outcome in stroke patients, we identified underlying neutrophil-driven mechanisms and explored therapeutic strategies to prevent the deleterious effects of pneumonia.

## RESULTS

### Stroke-associated pneumonia is associated with poor stroke outcome in ischemic stroke patients

To characterize the effect of stroke-associated pneumonia in acute ischemic stroke patients, we prospectively examined clinical outcome in the prospective NOFF-S cohort. Patients experiencing stroke-associated pneumonia (26 patients) during their hospital stay had similar age (70.0(61.5;79.5) vs. 70.0(60.0;78.3) years, respectively) and sex distribution (57.7% vs. 60.7%) as patients without pneumonia (326 patients), but more severe strokes at admission defined by the National Institute of Health Stroke Scale (NIHSS) score (12.0(9.5;15.0) vs. 4.0(2.0;7.0)) (Table 1). Stroke outcome at 90 days was worse in stroke patients with pneumonia than patients without pneumonia (modified Rankin Scale (mRS) score 2.0(1.0;4.0)) vs. 1.0(0.0;2.0), p=0.043) (Table 1). Pneumonia predicted poor stroke outcome at 90 days in univariate analyses (relative risk (RR) 3.2(1.0-9.9), p=0.044) and multivariable analyses adjusted for age and sex (3.2(1.0-9.9), p=0.045). RR only modestly decreased to 2.3(0.7-7.4) after age, sex, and NIHSS adjustment.

**Table 1.** Baseline characteristics and clinical outcome of ischemic stroke patients with and without stroke-associated pneumonia in the NOFF-S cohort.

| Variables | Stroke-associated pneumonia (n=26 patients) | No stroke-associated pneumonia (n=326 patients) | p |
| --- | --- | --- | --- |
| **Demographics** |  |  |  |
| Age | 70.0(61.5;79.5) | 70.0(60.0;78.3) | 0.784 |
| Male sex | 15 (57.7) | 198 (60.7) | 0.760 |
| **Stroke severity** |  |  |  |
| NIHSS at admission | 12.0(9.5;15.0) | 4.0(2.0;7.0) | <0.001 |
| NIHSS T1 | 11.0(6.3;14.8) | 3.0(1.0;5.0) | <0.001 |
| mRS T1 | 5.0(4.0;5.0) | 2.0(1.0;3.5) | <0.001 |
| Barthel Index T1 | 20.0 (5.0;40.0) | 85.0 (60.0;100.0) | <0.001 |
| MoCA T1 | 17.0(11.0;20.0) | 22.0(18.0;25.0) | 0.013 |
| NIHSS T2 | 10.0(5.0;13.0) | 2.0(1.0;4.0) | <0.001 |
| mRS T2 | 4.0(4.0;5.0) | 2.0(1.0;3.0) | <0.001 |
| Barthel Index T2 | 20.0 (10.0;42.5) | 95.0 (75.0;100.0) | <0.001 |
| MoCA T2 | 17.0(15.0;20.0) | 22.0(18.0;26.0) | 0.001 |
| **Cardiovascular risk factors** |  |  |  |
| Arterial hypertension | 17 (65.4) | 247 (76.2) | 0.216 |
| Diabetes mellitus | 7 (26.9) | 92 (28.4) | 0.873 |
| Dyslipidemia | 13 (50.0) | 163 (50.3) | 0.976 |
| Atrial fibrillation | 13 (50.0) | 69 (21.3) | <0.001 |
| Current smoking | 3 (15.0) | 75 (25.2) | 0.283 |
| BMI | 27.1(24.4;32.7) | 26.6(23.5;30.5) | 0.630 |
| **Cardiovascular diseases** |  |  |  |
| History of myocardial infarction | 2 (7.7) | 27 (8.3) | 0.909 |
| Coronary artery disease | 5 (19.2) | 60 (18.5) | 0.721 |
| Peripheral artery disease | 4 (8.2) | 13 (4.3) | 0.928 |
| **Interventions** |  |  | <0.001 |
| Thrombolysis | 3 (11.5) | 77 (23.8) |  |
| Thrombectomy | 6 (23.1) | 14 (4.3) |  |
| Thrombolysis plus thrombectomy | 6 (23.1) | 27 (8.3) |  |
| None | 11 (42.3) | 206 (63.6) |  |
| **Clinical Outcomes** |  |  |  |
| mRS T3 | 2.0(1.0;4.0) | 1.0(0.0;2.0) | 0.043 |
| NIHSS T3 | 1.0(1.0;1.0) | 1.0(0.0;1.0) | 0.144 |
| Barthel Index T3 | 100.0(90.0;100.0) | 100.0(100.0;100.0) | 0.154 |
| MoCA T3 | 24.0(23.0;24.0) | 25.0(22.0;27.0) | 0.368 |

Unless otherwise indicated, data are median(Q1;Q3) or numbers (%). P-values were
assessed by Mann-Whitney U tests for continuous data, categorical data by Chi-square or
Fisher's exact tests. T1: 24-<72 hours after admission, T2: 72-144 hours after admission,
T3: after 3 months.

### Bacterial pneumonia exacerbates post-ischemic neurological deficits, cerebral microvascular permeability, and thromboinflammation in mice

To explore the neuropathological correlates associated with stroke-associated pneumonia, we exposed mice to transient intraluminal MCAO and induced experimental pneumonia 72 hours thereafter by intratracheal *S. pneumoniae* instillation (time-line in Figure 1A). Experimental pneumonia decreased rectal temperature for up to 12 hours (Extended Data Fig. 1) and increased neurological deficits at 1 and 2 days post-pneumonia (dpp) (Fig. 1B). Pneumonia did not influence infarct volume at 2 dpp (i.e., 5 days post-MCAO) (Fig. 1C), but significantly increased brain edema (Fig. 1D) and IgG extravasation (Fig. 1E), which are markers of BBB breakdown. Pneumonia also increased intercellular adhesion molecule-1 (ICAM1) abundance on cerebral microvessels (Fig. 1F), increased brain CD45^+^ leukocyte infiltrates (Fig. 1G), increased the density of GP-Ibα^+^ microvascular thrombi (Fig. 1H), reduced the density of Iba1^+^ microglia (Fig. 1I), and increased the activation of Iba1^+^ microglia (evidenced by a reduced ramification index; Fig. 1J-L) in the previously ischemic brain tissue. Thus, stroke-associated pneumonia induced a robust surrogate of structural brain injury associated with the neurological deterioration.

**Fig 1.**
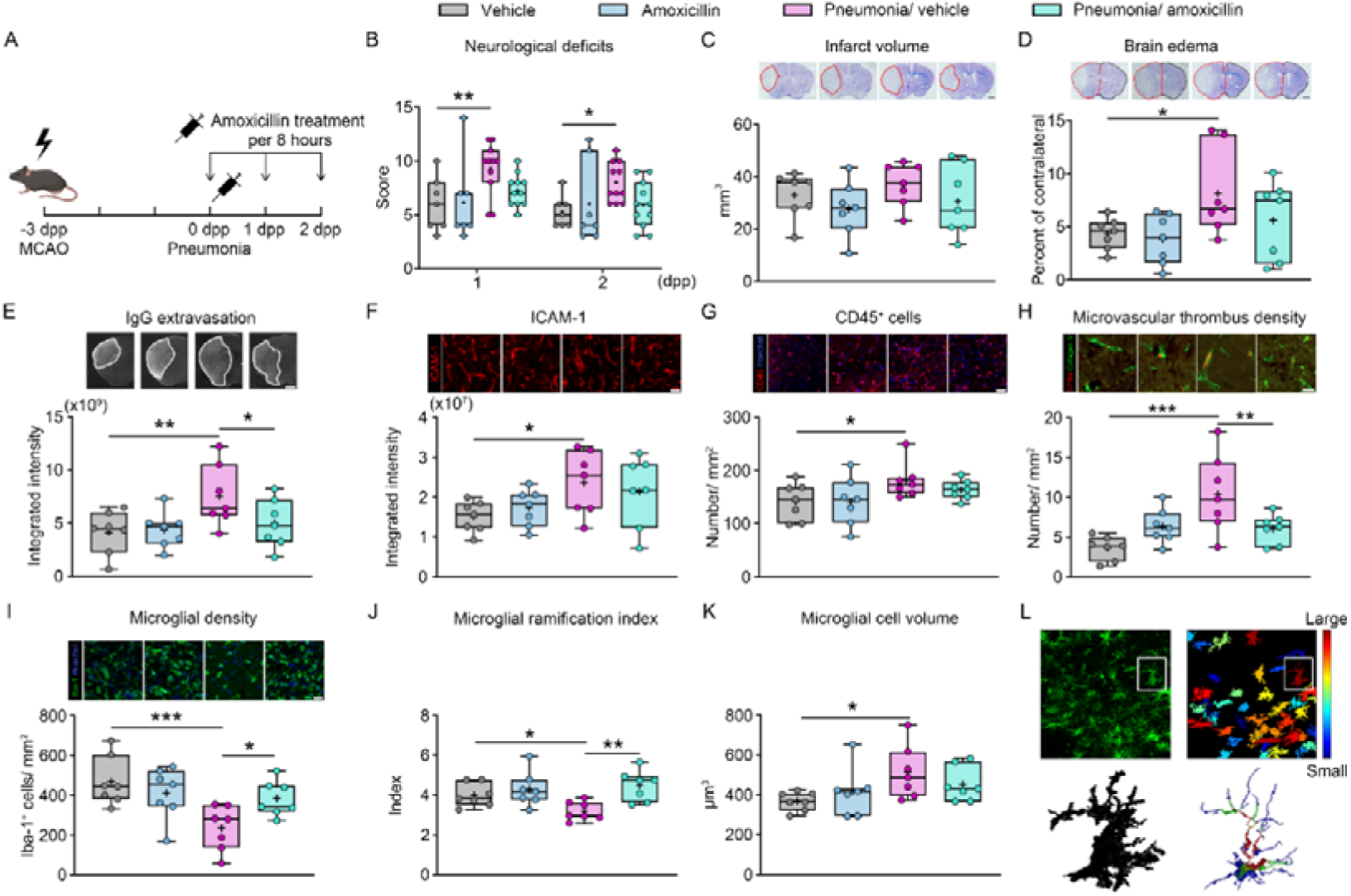
*S. pneumoniae* pneumonia increases neurological deficits, blood-brain barrier (BBB) breakdown, microvascular thrombus formation, and brain inflammatory responses post-stroke, which are incompletely reversed by amoxicillin. **(A)** Time-line of animal experiments, **(B)** neurological deficits assessed by a comprehensive behavioral score, **(C)** infarct volume, and **(D)** brain edema evaluated by cresyl violet staining, **(E)** IgG extravasation in ischemic brain tissue, **(F)** ICAM1 expression on ischemic microvessels, **(G)** brain infiltrated CD45^+^ leukocytes, **(H)** microvascular thrombus density in ischemic brain tissue, **(I-L)** density and activation of Iba1^+^ microglia in the ischemic striatum (i.e., the core of the middle cerebral artery (MCA) vascular territory), assessed by cell counting and morphology analysis following the segmentation, skeletonization, and reconstruction of Iba1^+^ cells of mice exposed to transient intraluminal MCAO, followed by no pneumonia or pneumonia induced by intratracheal *S. pneumoniae* instillation (1x10^8^ CFUs) at 3 days post-MCAO and vehicle or amoxicillin (15 mg/kg t.i.d., s.c.) treatment starting 3 hours thereafter. Mice were sacrificed at 2 days post-pneumonia (dpp; i.e., 5 days post-MCAO). Representative cresyl violet staining and immunohistochemistry images are shown. *p<0.05, **p<0.01, ***p<0.001 (n=7-11 mice/ group). Scale bars: 1 mm (in **(C-E)**); 50 μm (in **(F, G)**); 30 μm (in **(H, I)**).

### Antibiotic amoxicillin restores post-ischemic BBB permeability and microvascular thrombosis, but incompletely reverses the pneumonia-associated neurological deficits and microvascular inflammation

Antibiotic treatment with amoxicillin initiated 3 hours post-pneumonia reduced the rectal temperature decrease in MCAO mice (Extended Data Fig. 1). Neurological deficits post-pneumonia were not significantly reduced by amoxicillin (Fig. 1B). Amoxicillin did not significantly affect brain infarct volume at 2 dpp (Fig. 1C), but attenuated the pneumonia-associated BBB breakdown (Fig. 1D, E), but not ICAM1 levels on cerebral microvessels (Fig. 1F) or CD45^+^ leukocyte brain infiltrates (Fig. 1G). Amoxicillin reduced the GP-Ibα^+^ microvascular thrombus density in the previously ischemic tissue of MCAO/ pneumonia mice (Fig. 1H). Similarly, amoxicillin restored the density of Iba1^+^ microglia and reduced their activation (Fig. 1I-L). Hence, antibiotic treatment did not fully reverse the neurological worsening and histopathological injury in stroke-associated pneumonia mice, in line with randomized controlled trials in stroke patients, in which antibiotics failed to enhance stroke outcome ^11,12^.

### Amoxicillin reverses the pneumonia-associated neutrophil responses in the blood, but not ischemic brain

To elucidate mechanisms underlying the pneumonia-associated brain damage, we performed flow cytometry of leukocytes obtained from peripheral blood and ischemic brain tissue at 2 dpp using the sorting strategy provided in Extended Data Fig. 2. Pneumonia did not significantly alter the overall number of CD45^+^ leukocytes in peripheral blood, but significantly increased the number and activation of Ly6G^+^ neutrophils and CD8^+^ T cell activation (Fig. 2). Amoxicillin reversed the elevated neutrophil number and activation and reduced the CD4^+^ and CD8^+^ T cell number in the blood of MCAO/ pneumonia mice (Fig. 2).

**Fig 2.**
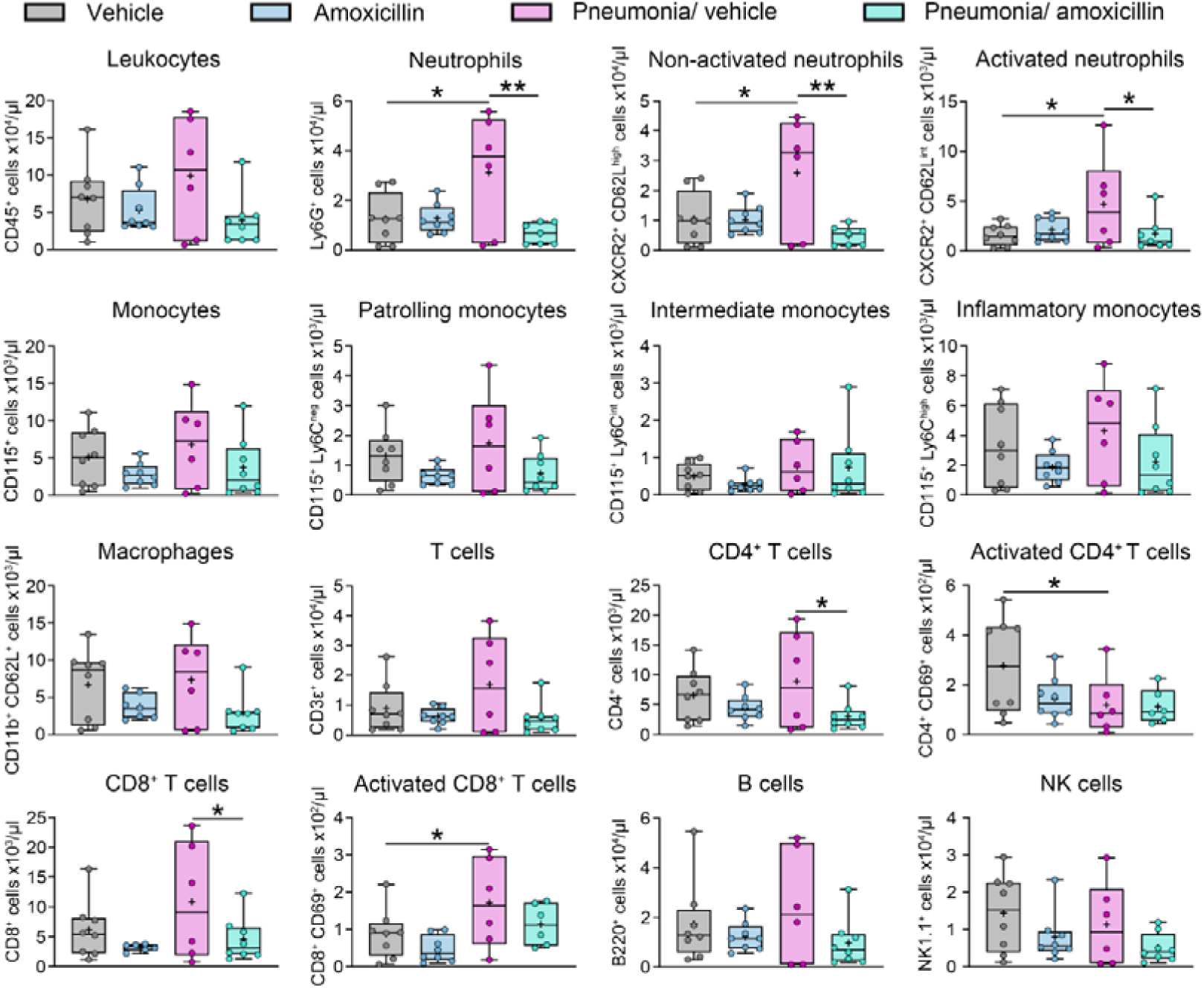
Pneumonia increases the number and activation of neutrophils in peripheral blood, which is reversed by amoxicillin. Number of CD45^+^ leukocytes, Ly6G^+^ neutrophils, Ly6G^+^ CXCR2^+^ CD62L^high^ non-activated neutrophils, Ly6G^+^ CXCR2^+^ CD62L^int^ activated neutrophils, CD115^+^ monocytes, CD115^+^ Ly6C^neg^ patrolling monocytes, CD115^+^ Ly6C^int^ intermediate monocytes, CD115^+^ Ly6C^high^ inflammatory monocytes, CD115^-^ CD11b^+^ CD62L^+^ macrophages, CD3ε^+^ T cells, CD4^+^ T cells, CD4^+^ CD69^+^ activated T cells, CD8^+^ T cells, CD8^+^ CD69^+^ activated T cells, B220^+^ B cells, and NK1.1^+^ NK cells in peripheral blood assessed by flow cytometry of mice exposed to transient intraluminal MCAO, followed by no pneumonia or *S. pneumoniae* pneumonia at 3 days post-MCAO and vehicle or amoxicillin treatment (15 mg/kg t.i.d., s.c.) starting 3 hours later. Animals were sacrificed at 2 dpp. Note that besides neutrophils, CD8^+^ activated T cells were also increased by pneumonia. Amoxicillin reduced CD8^+^ T cell numbers in peripheral blood of pneumonic mice. *p<0.05, **p<0.01 (n=6-8 mice/ group).

Pneumonia also increased the number of brain-infiltrated CD45^+^ leukocytes, including Ly6G^+^ neutrophils, CD115^+^ monocytes (particularly patrolling and intermediate monocytes), CD3ε^+^ T cells, B220^+^ B cells, and NK1.1^+^ NK cells in the previously ischemic brain tissue (Fig. 3). Of note, immune cell numbers in the brain, including neutrophil numbers, were not influenced by amoxicillin (Fig. 3).

**Fig 3.**
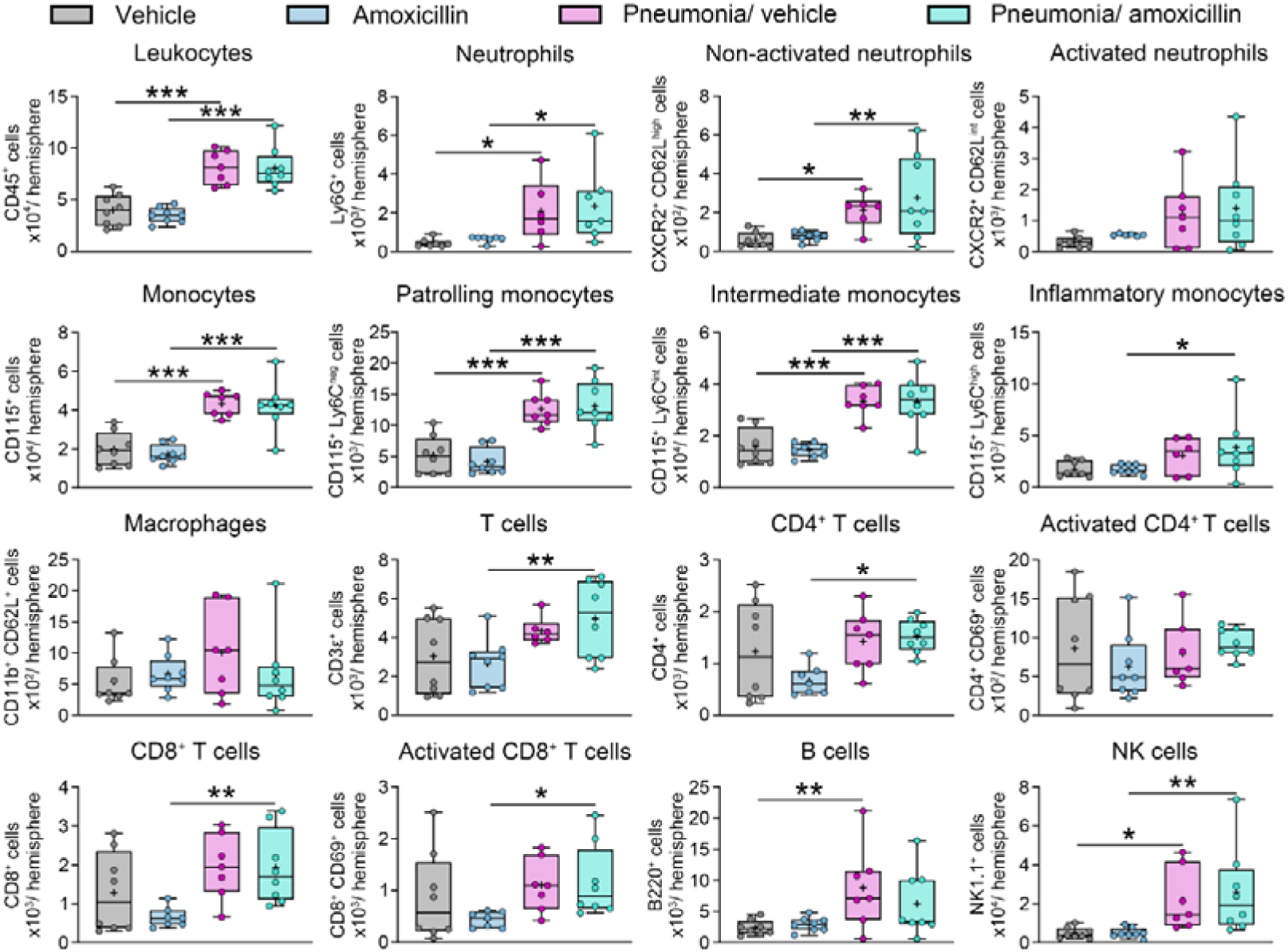
Pneumonia increases the brain infiltration of neutrophils, monocytes, T cells, B cells, and NK cells, which are not reversed by amoxicillin. Number of infiltrated CD45^+^ leukocytes, Ly6G^+^ neutrophils, Ly6G^+^ CXCR2^+^ CD62L^high^ non-activated neutrophils, Ly6G^+^ CXCR2^+^ CD62L^int^ activated neutrophils, CD115^+^ monocytes, CD115^+^ Ly6C^neg^ patrolling monocytes, CD115^+^ Ly6C^int^ intermediate monocytes, CD115^+^ Ly6C^high^ inflammatory monocytes, CD115^-^ CD11b^+^ CD62L^+^ macrophages, CD3ε^+^ T cells, CD4^+^ T cells, CD4^+^ CD69^+^ activated T cells, CD8^+^ T cells, CD8^+^ CD69^+^ activated T cells, B220^+^ B cells, and NK1.1^+^ NK cells in the ischemic brain assessed by flow cytometry of mice exposed to transient intraluminal MCAO, followed by sham-exposure or *S. pneumoniae* pneumonia at 3 days post-MCAO and vehicle or amoxicillin treatment (15 mg/kg t.i.d., s.c.) starting 3 hours later. Animals were sacrificed at 2 dpp. *p<0.05, **p<0.01, ***p<0.001 (n=7-8 mice/ group).

### Pneumonia induces a degranulation, platelet activation, and NETosis signature in peripheral blood neutrophils

To further explore the role of neutrophils in stroke-associated pneumonia, we collected blood samples of MCAO mice receiving vehicle or MCAO/ pneumonia mice receiving vehicle or amoxicillin at 2 dpp (that is, 5 days post-MCAO), from which neutrophils were isolated for proteome analysis. In total, 1058 proteins were detected in neutrophil samples. In a comparison between the MCAO/ pneumonia/ vehicle and the MCAO only/ vehicle group, 35 proteins with p-values less than 0.05 and absolute log_2_(fold change) larger than 0.263 were identified as differentially expressed, was shown by a heatmap of their relative expression levels (Fig. 4A). Among these 35 proteins, 26 were upregulated and 9 were downregulated by pneumonia. Among the proteins increased by pneumonia, 7 were related to neutrophil degranulation (Cct8, Cotl1, Mvp, Arl8a, Tollip, Cap1, Aldoa) and 5 related to platelet activation (Cap1, Aldoa, Actn4, Gnai3, Anxa5) (Fig. 4B). In a comparison between the MCAO/ pneumonia/ amoxicillin and MCAO/ pneumonia/ vehicle groups, 64 proteins were differentially expressed with p-values less than 0.05 and absolute log_2_(fold change) larger than 0.263, with 34 downregulated and 30 upregulated proteins (Fig. 4C). Among the 34 downregulated proteins, 5 were related to neutrophil degranulation (Cand1, Psma2, Cotl1, Arl8a, Cap1), 5 to platelet activation (Cap1, Anxa5, Csk, Actn4, Gnai3), and one to cell and vascular wall adhesion (Bsg) (Fig. 4D). Besides, 2 proteins related to neutrophil degranulation (Ctsg, Psma5) and 2 associated with platelet activation (Itih3, Col1a2) were increased by amoxicillin (Fig. 4D). 12 proteins showed opposite regulation in the two comparisons (MCAO/ pneumonia/ amoxicillin vs. MCAO/ pneumonia/ vehicle; MCAO/ pneumonia/ vehicle vs. MCAO only/ vehicle) (Fig. 4E), of which 9 proteins were increased by pneumonia and decreased by amoxicillin. Among these 9 proteins, 3 were related to neutrophil degranulation (Arl8a, Cap1, Cotl1) and 4 were related to platelet activation (Cap1, Gnai3, Anxa5, Actn4).

**Fig. 4.**
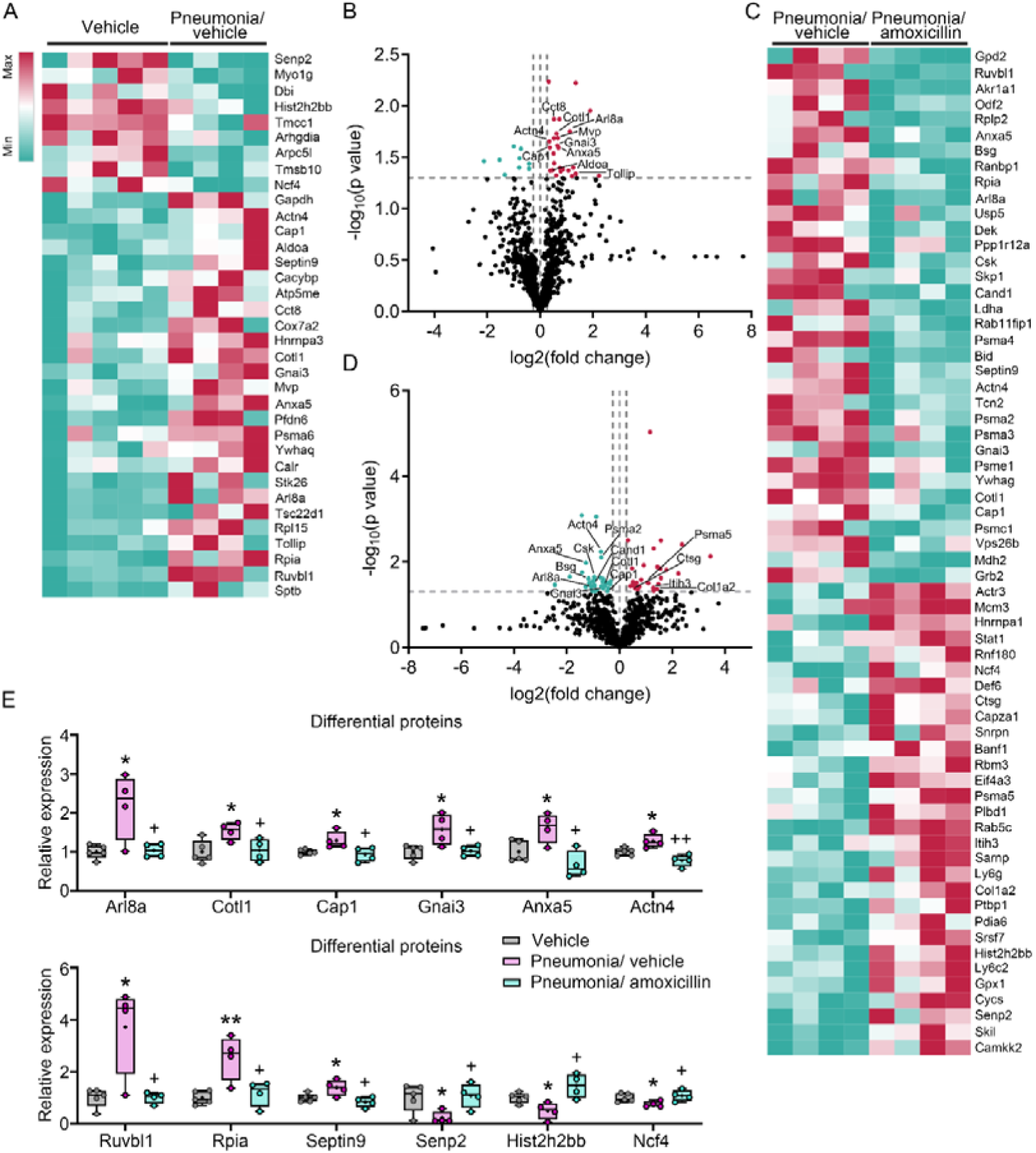
Pneumonia induces a neutrophil degranulation and NETosis signature that is reversed by amoxicillin. **(A)** Heatmap and **(B)** volcano plot of differentially expressed proteins in peripheral blood neutrophils of MCAO mice exposed to *S. pneumoniae* pneumonia at 3 days post-MCAO treated with vehicle (s.c.) compared with MCAO mice without pneumonia treated with vehicle that were sacrificed at 2 dpp. **(C)** Heatmap and **(D)** volcano plot of differentially expressed proteins of peripheral blood neutrophils of MCAO/ pneumonia mice treated with amoxicillin (15 mg/kg t.i.d., s.c.) starting 3 hours post-pneumonia compared with MCAO/ pneumonia mice treated with vehicle that were sacrificed at 2 dpp. **(E)** Proteins significantly regulated in the three conditions in opposing ways. Note that besides proteins involved in neutrophil degranulation (Arl8a, Cotl1, Cap1) and platelet activation (Cap1, Gnai3, Anxa5, Actn4), the histone 2B family protein Hist2h2bb and Ruvbl1 protein that promotes chromatin decondensation, histone modification, and chromatin remodeling were identified. *p<0.05, **p<0.01 for MCAO/ pneumonia/ vehicle compared with MCAO only/ vehicle; ^+^p<0.05, ^++^p<0.01 for MCAO/ pneumonia/ amoxicillin compared with MCAO/ pneumonia/ vehicle (n=4-5 mice/ group).

We also found two proteins, Ruvbl1 and Hist2h2bb, which were oppositely regulated in the three groups. Ruvbl1 (UniProt #P60122) is a RuvB-like-1 protein that promotes chromatin decondensation, histone modification, and ATP-dependent chromatin remodeling, while Hist2h2bb (UniProt #P62806) is a histone 2B family protein. Ruvbl1 expression was increased and Hist2h2bb expression was decreased in the pneumonia/ vehicle compared to the MCAO only/ vehicle group, while Ruvbl1 was decreased and Hist2h2bb was increased in the pneumonia/ amoxicillin compared to pneumonia/ vehicle group (Fig. 4E). We interpreted the Rubbl1 elevation post-pneumonia as possible indicator of ongoing NETosis and the Hist2h2bb decrease as indicator of prior histone protein release.

### Neutrophils play a central role in brain injury associated with pneumonia

To elucidate the role of neutrophils in the pneumonia-associated brain injury, we depleted neutrophils by administering a monoclonal anti-Ly6G (1A8) antibody at 1 day before and 1 day after pneumonia (i.e., at 2 and 4 days post-MCAO) (time-line see Extended Data Fig. 3A). Neutrophil depletion at these subacute time-points did not influence infarct volume, brain edema, IgG extravasation, and microvascular thrombus density at 2 dpp (that is, 5 days post-MCAO) in MCAO only mice, but reduced infarct volume, brain edema, IgG extravasation, and microvascular thrombus density in MCAO/ pneumonia mice (Fig. 5A-D). Neurological deficits, ICAM1 expression on cerebral microvessels and the density of brain-infiltrating CD45^+^ leukocytes were not influenced by neutrophil depletion in MCAO only or MCAO/ pneumonia mice (Extended Data Fig. 3B; Extended Data Fig. 4A, B). Brain hemorrhage formation was not influenced by neutrophil depletion, neither with respect to hemorrhage volume nor hemorrhage incidence (Extended Data Fig. 4C, D). These findings demonstrate a central role of neutrophils in the histopathological injury associated with pneumonia.

**Fig. 5.**
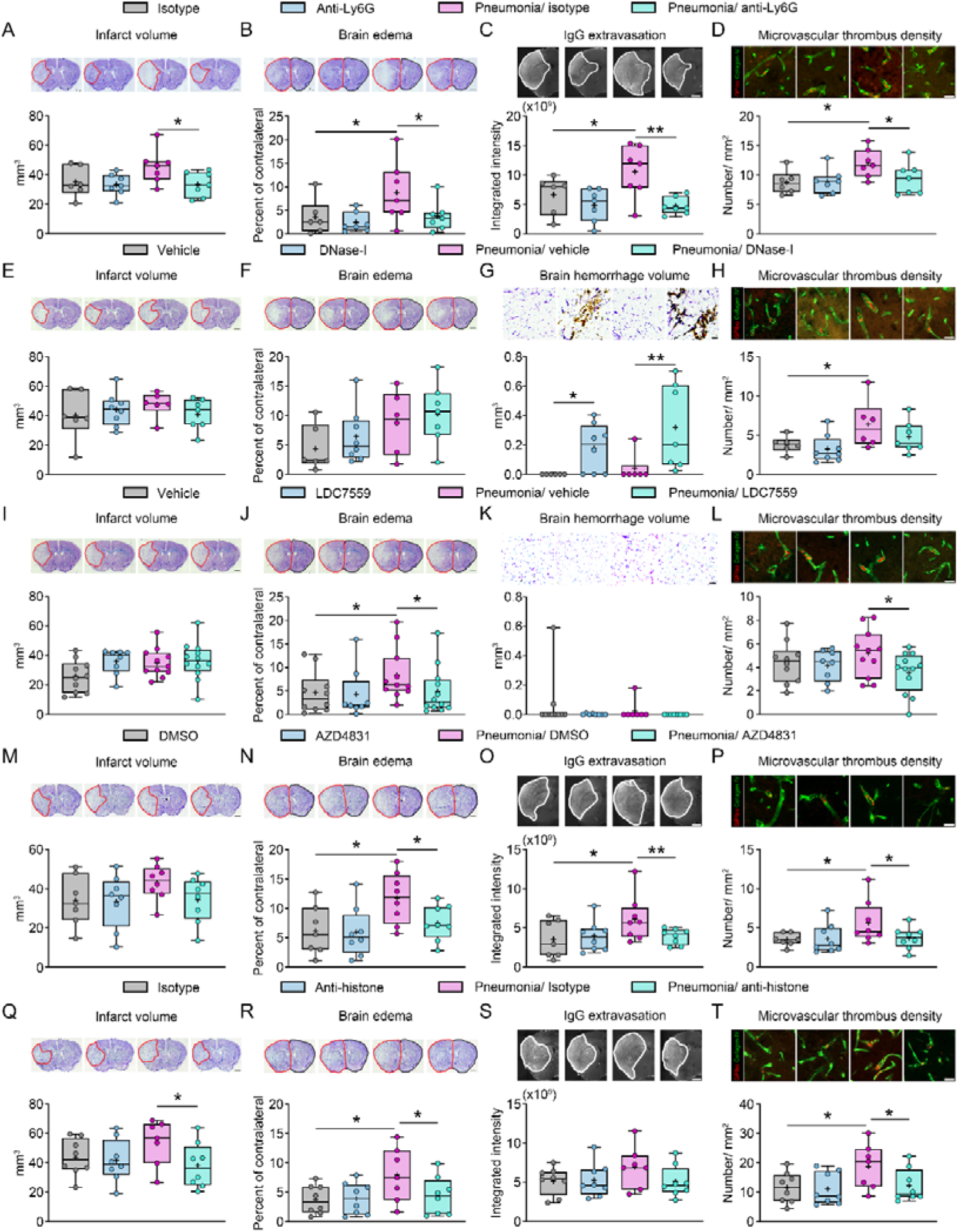
Antibody-mediated neutrophil depletion, NET formation blockade, MPO inhibition, and histone neutralization reverse the early neuropathological sequelae associated with pneumonia. **(A, E, I, M, Q)** Infarct volume, **(B, F, J, N, R)** brain edema, **(C, O, S)** IgG extravasation, **(D, H, L, P, T)** GP-Ibα^+^ thrombus density (in red) on collagen-IV^+^ cerebral microvessels (in green), and **(G, K)** brain hemorrhage volume in the previously ischemic brain tissue of MCAO mice without pneumonia or MCAO mice exposed to *S. pneumoniae* pneumonia at 3 days post-MCAO, which were treated with **(A-D)** isotype antibody or anti-Ly6G antibody (1A8; 200 µg i.p.), which near-completely depletes neutrophils ^15^, 1 day before and 1 day after pneumonia (i.e., at 2 and 4 days post-MCAO), **(E-H)** vehicle or DNase-I (10 µg i.v. and 50 µg i.p. 30 min before pneumonia, followed by 50 µg b.i.d.), which degrades NETs ^19^, **(I-L)** vehicle or the gasdermin-D inhibitor LDC7559 (10 mg/kg i.p.), which prevents NET formation ^19^, 30 min before pneumonia and 1 day after pneumonia, **(M-P)** vehicle or the MPO inhibitor AZD4831 (10 µmol/kg via oral gavage) 2 hours before and 1 day after pneumonia, or **(Q-T)** isotype IgG or neutralizing anti-histone antibody (BWA3; 20 µg i.p.) 30 min before and 1 day after pneumonia, followed by animal sacrifice at 2 dpp (that is, 5 days post-MCAO). Representative cresyl violet and immunohistochemistry images are shown. *p<0.05, **p<0.01 (n=6-12 mice/ group). Scale bars: 1 mm (in **(A-C**, **E, F**, **I, J**, **M-O**, **Q-S)**); 30 µm (in **(D**, **H**, **L**, **P**, **T)**); 50 μm (in **(G**, **K)**).

### NET degradation by DNase-I fails to alleviate the neuropathological consequences of pneumonia, but promotes brain hemorrhages

Considering that neutrophils in peripheral blood carried a protein signature associated with neutrophil degranulation and NETosis, we next asked if the deleterious effects of pneumonia on ischemic injury were attributed to NETs, which are released by neutrophils upon severe cellular stress ^17^. To clarify this question, we evaluated the effects of DNase-I administration starting 30 minutes before pneumonia (i.e., 3 days post-MCAO) (Extended Data Fig. 3C), which as we have previously shown degrades NETs ^19^. DNase-I did not influence neurological deficits, infarct volume, brain edema, IgG extravasation, and microvascular thrombosis at 2 dpp (5 days post-MCAO) in MCAO only or MCAO/ pneumonia mice (Extended Data Fig. 3B; Fig. 5E, F, H; Extended Data Fig. 4E). Similar to neutrophil depletion, DNase-I did not alter microvascular ICAM1 expression and brain-infiltrating CD45^+^ leukocyte density (Extended Data Fig. 4F, G). The lack of recovery-promoting effects was possibly due to the exacerbation of brain hemorrhage volume and incidence, which were increased by DNase-I in MCAO and MCAO/ pneumonia mice (Fig. 5G; Extended Data Fig. 4H). Hence, due to the risk of brain bleeding, DNase-I administration should be considered in the post-acute stroke phase with care.

### NET formation blockade by gasdermin-D inhibitor prevents brain edema and microvascular thrombosis without increasing brain hemorrhages

Since NET degradation by DNase-I induced brain hemorrhages, we asked if a more subtle intervention that does not entirely eliminate preexisting NETs may be able to reverse the consequences of pneumonia. Thus, we evaluated the effects of a NET formation blocker, the pharmacological gasdermin-D inhibitor LDC7559 (HY-111674; MedChemExpress, Monmouth Junction, NJ, U.S.A.) ^21^, which we administered 30 min before pneumonia (Extended Data Fig. 3E). LDC7559 delivery reversed the increased levels of CitH3^+^ DNA, which is specific for NETs ^19^, in peripheral blood plasma (Extended Data Fig. 5). Although LDC7559 administered at this delayed time-point did not influence neurological deficits (Extended Data Fig. 3F) and infarct volume (Fig. 5I), it restored pneumonia-induced BBB integrity and reduced microvascular thrombosis (Fig. 5J, L; Extended Data Fig. 4I) at 2 dpp. ICAM1 expression on cerebral microvessels and brain-infiltrating CD45^+^ leukocyte density were not influenced by LDC7559 (Extended Data Fig. 4J, K). Of note, NET formation blockade by LDC7559 did not increase brain hemorrhage volume or incidence (Fig. 5K; Extended Data Fig. 4L). Hence, LDC57559 reversed the structural consequences of pneumonia in the subacute stroke phase.

### Myeloperoxidase inhibition restores the early neuropathological, but not neurological consequences of pneumonia

Once released by neutrophils, NETs aggregate with neutrophil-derived proteins, namely myeloperoxidase (MPO) and histones including histone H4, which induce cell injury under ischemia/reperfusion conditions ^22,23^. To clarify the contribution of MPO to pneumonia-associated brain injury, we administered the pharmacological inhibitor AZD4831 (HY-145581; MedChemExpress) ^24^ 2 hours before pneumonia (Extended Data Fig. 3G). Plasma MPO activity doubled upon pneumonia, and MPO inhibition by AZD4831 reversed it (Extended Data Fig. 6). AZD4831 did not influence neurological deficits (Extended Data Fig. 3H) and infarct volume (Fig. 5M) at 2 dpp, but reestablished pneumonia-induced BBB integrity, indicated by brain edema and IgG extravasation (Fig. 5N, O), and reduced microvascular thrombosis (Fig. 5P). ICAM1 expression on cerebral microvessels, brain CD45^+^ leukocyte infiltrates and intracerebral hemorrhages were not altered by AZD4831 (Extended Data Fig. 4M-P). Thus, AZD4831 reversed the structural, but not functional neurological consequences of pneumonia.

### Histone blockade prevents the early neurological and neuropathological consequences of pneumonia

To neutralize histones, we administered the monoclonal anti-histone-H4/H2A antibody BWA3 (Creative Biolabs, Shirley, NY, USA) ^25^ 30 minutes before and 1 day after pneumonia (Extended Data Fig. 3I). Histone blockade reduced neurological deficits (Extended Data Fig. 3J) and infarct volume (Fig. 5Q) at 2 dpp in MCAO/ pneumonia mice, reversed the pneumonia-associated BBB breakdown (Fig. 5R, S) and reduced microvascular thrombosis (Fig. 5T). Microvascular ICAM1 expression, brain CD45^+^ leukocyte infiltrates, and intracerebral hemorrhages were not influenced by histone blockade (Extended Data Fig. 4Q-T). Taken together, histone blockade reversed the neurological and neuropathological consequences of pneumonia in the early stroke recovery phase.

### Pneumonia worsens long-term neurological and histopathological stroke outcome

To elucidate the long-term consequences of pneumonia for functional and structural stroke outcome, we exposed MCAO mice to *S. pneumoniae* pneumonia at 3 days post-MCAO and examined neurological recovery over 56 dpp (time-line see Fig. 6A). Pneumonia significantly exaggerated behavioral deficits assessed by the neurological score, tight rope and Rotarod tests, which did not fully recover over 56 dpp (Fig. 6B-D). Of note, brain atrophy at 56 dpp was significantly increased by pneumonia (Fig. 6E), indicating compromised structural brain remodeling in stroke-associated pneumonia mice.

**Fig. 6.**
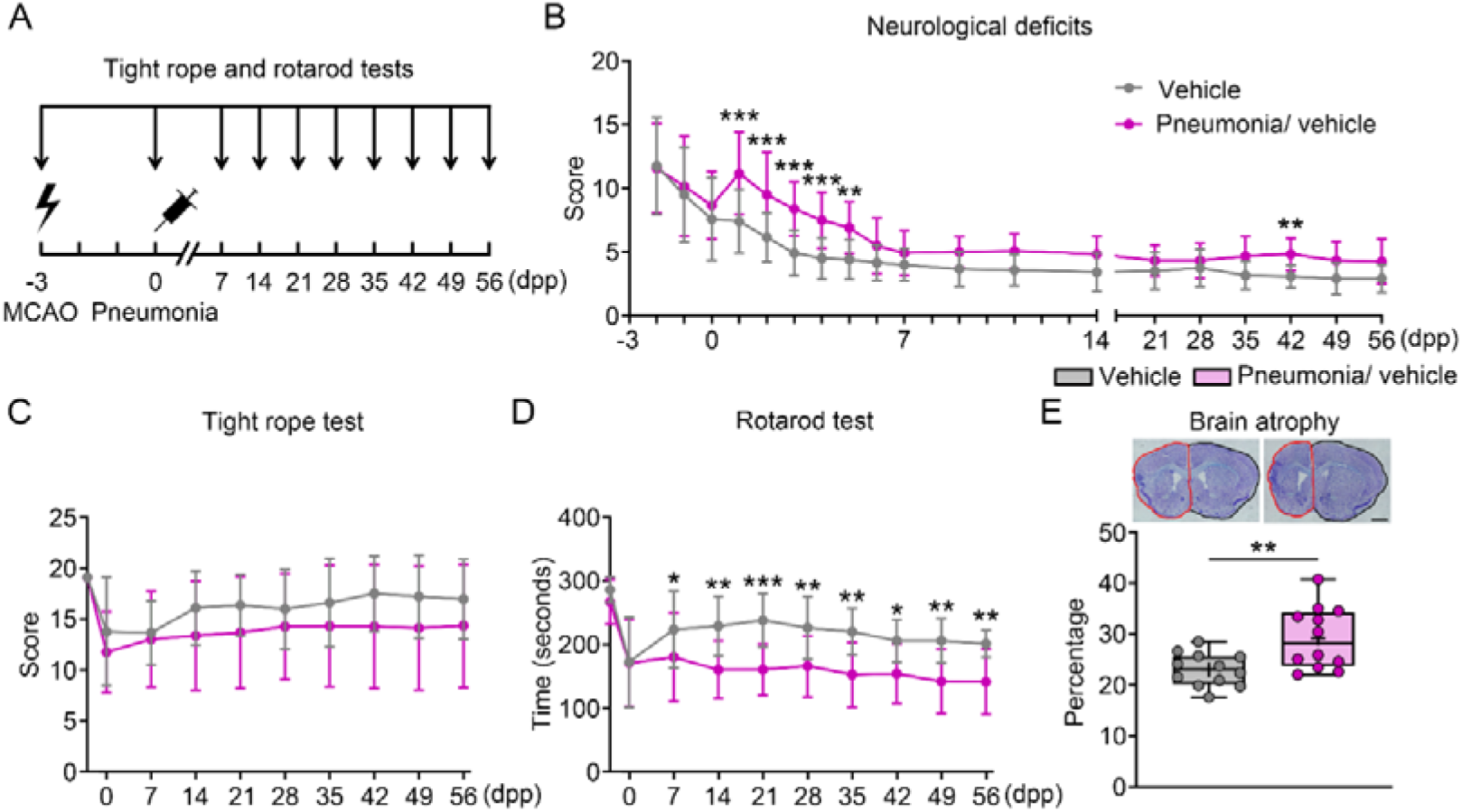
Pneumonia compromises long-term functional and structural stroke outcome. **(A)** Time-line of animal experiments, **(B-D)** functional deficits evaluated using the neurological score, tight rope and RotaRod tests, and **(E)** brain atrophy of mice exposed to transient intraluminal MCAO, followed by no pneumonia or *S. pneumoniae* pneumonia at 3 days post-MCAO. Animals were sacrificed at 56 dpp. Representative cresyl violet stainings are depicted. *p<0.05, **p<0.01, ***p<0.001 (n=12/ group). Scale bars: 1 mm (in **(E)**).

### NET formation inhibition during pneumonia does not influence long-term stroke recovery

Considering its effects on BBB integrity and microvascular thrombus formation, we next asked if NET formation blockade using LDC7559 promotes long-term stroke recovery in MCAO/ pneumonia mice. LDC7559 treatment on occasion of pneumonia (that is, 3 days after MCAO; time-line in Extended Data Fig. 7A) did not enhance functional and structural stroke outcome assessed by the neurological score, tight rope, RotaRod, open field and novel object recognition tests, brain atrophy, and microglial density and activation over up to 56 dpp (Extended Data Fig. 7B-L). In contrast, LDC7559 administered immediately after MCAO (time-line in Extended Data Fig. 8A) significantly reduced behavioral deficits in the neurological score and tight rope tests, increased spontaneous motor activity in open field tests, increased novel object discrimination in object recognition tests, reduced brain trophy, and reduced microglial activation (Extended Data Fig. 8B-L). Our results indicate that NET formation blockade was effective only at early stages before NET release has taken place.

### Histone blockade during pneumonia induces long-term functional and structural stroke recovery

Considering its effects on the functional and structural sequelae of pneumonia in the early stroke recovery phase, we finally asked if histone blockade using the BWA3 antibody enhances long-term stroke recovery post-pneumonia. Histone blockade during pneumonia (that is, 3 days after the stroke; time-line in Fig. 7A) reduced behavioral deficits assessed by neurological score and tight rope tests (Fig. 7B, C), increased spontaneous motor activity in open field tests (Fig. 7D) and novel object recognition in object recognition tests (Fig. 7E), and reduced brain atrophy (Fig. 7F) evolving over up to 56 dpp. As possible underlying mechanism, histone blockade also increased microglial density (Fig. 7G) and reduced microglial activation assessed by ramification index and cell volume (Fig. 7H-J). Thus, histone blockade persistently improved long-term neurological and histopathological outcome even when administered 3 days after stroke.

**Fig. 7.**
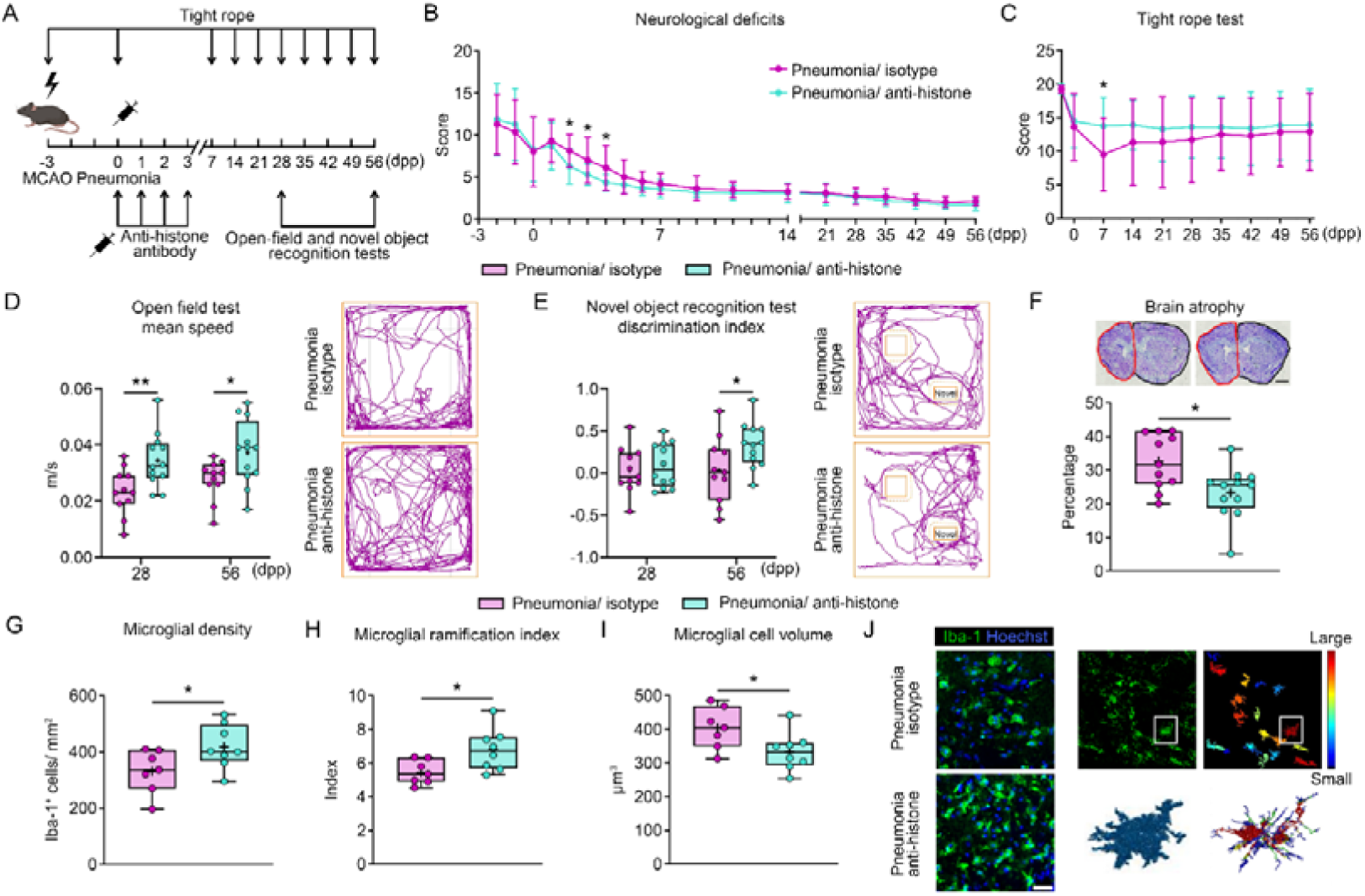
Histone blockade promotes long-term neurological recovery and prevents brain atrophy in mice with stroke-associated pneumonia, when administered in the post-acute stroke phase. **(A)** Time-line of animal experiments, **(B, C)** functional deficits evaluated by the neurological score and tight rope tests, **(D)** spontaneous motor activity in open field tests (tracking paths of representative animals at 56 dpp are shown), **(E)** novel object recognition in object recognition tests (tracking paths of representative animals at 56 dpp are also shown), **(F)** brain atrophy, assessed by cresyl violet staining, and **(G-J)** density and activation of Iba1^+^ microglia in the ischemic striatum, assessed by cell counting and morphology analysis following microglia segmentation, skeletonization and reconstruction, of MCAO mice exposed to *S. pneumoniae* pneumonia, which were treated with isotype IgG or neutralizing anti-histone antibody (BWA3; 20 µg i.p.) starting 30 min before pneumonia, followed by animal sacrifice at 56 dpp (behavioral and histochemical studies in **(B-F)**) or 2 dpp (microglial density and morphology studies in **(G-J)**). Representative samples in **(F)** and **(J)** are depicted. *p<0.05, **p<0.01 (n=11-12 mice/ group (in **(B-F)**); n=7-8 mice/ group (in **(G-J)**)). Scale bar: 1 mm (in **(F)**); 30 μm (in **(J)**).

## DISCUSSION

Using complementary analyses in ischemic stroke patients and in mice subjected to middle cerebral artery occlusion, we demonstrate that stroke-associated pneumonia markedly worsens stroke outcome. In mice, this process is neutrophil-dependent. It involves BBB breakdown, cerebral microvascular thrombosis, and progressive brain atrophy. Antibiotic treatment with amoxicillin only partially ameliorated the pneumonia-associated neurological deficits, indicating that infection control alone is insufficient to prevent secondary brain damage. In contrast, extracellular histone neutralization improved the long-term neurological and neuropathological consequences of pneumonia. Of note, NET degradation, blockade or MPO inhibition provided no lasting functional and structural benefit. Our results establish histone neutralization as a therapeutic strategy that promotes recovery in the post-acute stroke phase.

Microvascular permeability and thrombosis are well-established features of bacterial pneumonia and sepsis ^26,27^. Infection-remote microvascular thrombosis is known from severe acute respiratory syndrome-corona virus-2 infection (SARS-CoV-2), particularly in the heart and less commonly the brain ^28,29^. Microvascular thrombosis in remote organs is rare in *S. pneumoniae* pneumonia, except for hemolytic uremic syndrome, a fulminant condition characterized by severe thrombocytopenia and hemolytic anemia ^30^. To our knowledge, BBB breakdown and cerebral microvascular thrombosis have not yet been reported in *S. pneumoniae* pneumonia, neither in stroke-associated pneumonia nor pneumonia in other contexts.

Neutrophils are key components of dynamic capillary stalls that impede the reopening of reperfused blood vessels, promoting microvascular thrombosis and ischemic brain damage ^31,32^. Given that neutrophils travel from the infected lungs to cerebral vessels, we hypothesized neutrophils mediate the exacerbated brain injury observed in stroke-associated pneumonia mice. Indeed, anti-Ly6G antibody-mediated neutrophil depletion reversed the detrimental effects of pneumonia on BBB breakdown and microvascular thrombosis, and reduced ischemic brain injury. However, neutrophils are central for bacterial clearance ^33^, and post-stroke immunodepression hampers adaptive immunity ^19^. Thus, neutrophil depletion is not a viable option in stroke patients. Since antibiotic treatment with amoxicillin failed to reduce brain neutrophil infiltration in our model, we sought alternative strategies to mitigate the harmful consequences of pneumonia on the ischemic brain.

Proteome analysis showed that stroke-associated pneumonia induces a degranulation, platelet activation, and NETosis signature in mouse neutrophils, which was partially reversed by amoxicillin. Microvascular thrombi are heavily enriched with neutrophil-derived DNA NETs ^34,35^, which promote platelet aggregation and thrombus formation ^34,36^, contributing to reperfusion resistance and ischemic damage after stroke ^36,37^. Given this, we investigated whether removing NETs could reverse the deleterious effects of stroke-associated pneumonia. Contrary to our hypothesis, NET depletion by DNase-I did not alleviate the pneumonia-associated deterioration of stroke damage. Instead, it increased hemorrhagic transformation of brain infarcts. These data suggest that NETs play a crucial role in sealing ischemic blood vessels, as previously proposed by us for MCAO conditions not associated with pneumonia ^21^.

Three randomized controlled trials currently assess the effects of intravenously administered DNase-I in ischemic stroke patients (clinicaltrials.gov/study/NCT05203224, NCT05880524, NCT04785066). These studies should carefully monitor signs of hemorrhagic transformation after DNase-I treatment.

To inhibit NET formation, we utilized the pharmacological gasdermin-D blocker LDC7559, which we previously showed to reduce blood NET levels in ischemic stroke mice ^19^. Unlike DNase-I, this blockade did not increase hemorrhagic transformation. While LDC7559 reduced BBB leakage and microvascular thrombosis in stroke-associated pneumonia mice, it only enhanced long-term neurological recovery and progressive brain atrophy when administered immediately after reperfusion. Our findings confirm that post-ischemic NET formation is a rapid process ^19^ requiring early intervention to improve stroke outcome ^36^.

Our next target, the NET-bound peroxidase MPO, activates a large number of proteases, including matrix metalloproteases (MMPs), via its product HOCl/OCl^−^ through conversion of cysteine into thiol residues ^38,39^. In case of MMPs, this conversion facilitates the cleavage of pro-MMPs to MMPs including MMP-9 ^38,40^, which contributes to BBB breakdown and brain injury after tissue-plasminogen activator-induced thrombolysis in embolic mouse stroke models ^41^. Due to its diffusible nature, HOCl/OCl^−^ was a strong candidate to mediate BBB breakdown and thrombosis in infection-remote vascular beds ^39^. Although MPO deactivation by AZD4831 antagonized the pneumonia-associated BBB breakdown and microvascular thrombosis, it did not restore neurological deficits, when administered during pneumonia. Pharmacological MPO inhibition has previously reduced neurological deficits, brain edema, and ischemic injury when initiated immediately after MCAO in mice ^42^.

In subsequent studies, extracellular histones emerged as a highly promising long-window therapeutic target. Indeed, histone neutralization during pneumonia with a monoclonal antibody improved neurological recovery in stroke-associated pneumonia mice and reversed BBB breakdown, microvascular thrombosis, and progressive brain atrophy, when administered as late as 72 hours after stroke. Consequentially, histone neutralization enhanced long-term functional and structural outcome. Extracellular histones, most notably histone-H4, are key NET components that induce smooth muscle cell damage in mouse atherosclerosis models ^23^. Injured smooth muscle cells were found to attract neutrophils, promoting the ejection of NETs containing nuclear proteins including histone-H4 ^23^. This reiterated process destabilizes atherosclerotic plaques, whereas histone-H4 neutralization stabilizes atherosclerotic plaques ^23^. To our knowledge, this is the first demonstration that targeting extracellular histones confers brain-protective effects in ischemic stroke.

Given the repeated clinical failure of neurorestorative therapies applied broadly across stroke populations ^5,7,43^, our findings advocate a shift toward mechanism-based, patient-tailored interventions. We propose that targeting neutrophil-driven inflammatory injury in patients with exaggerated immune responses—such as those with stroke-associated infections—represents a promising strategy to improve outcome. Due to its long time-window, extracellular histone blockade offers a clinically feasible intervention. Small-molecule inhibitors, inhibitory peptides, and neutralizing antibodies for histone-H4 are currently in development ^44^, underscoring the translational relevance of our work. We predict that moving away from a one-size-fits-all approach toward targeted restorative therapies for selected patient populations—such as those with stroke-associated infections—could enable meaningful breakthroughs in stroke recovery.

## MATERIALS AND METHODS

### Implications of stroke-associated pneumonia in acute ischemic stroke patients

In the Neutrophils: Origin, Fate & Function Stroke (NOFF-S) study recruited via the German Research Foundation (DFG) Collaborative Research Center (CRC) TRR332 “Neutrophils: origin, fate, and function” (https://www.neutrophils.de, subproject C6) (DRKS00030825), patients with first-ever acute ischemic stroke ≥18 years, who were admitted to the Stroke Unit of the University Hospital Essen ≤48 hours after symptom onset have prospectively been recruited since October 2022. The current data analysis of NOFF-S includes patients included until August 21, 2025. Within 24-<72 hours after admission (examination T1), extensive clinical examinations, structured interviews and blood laboratory tests were performed by a certified study clinician to determine stroke outcome and history of vascular risk factors and diseases. Examinations were repeated 72-144 hours after admission (T2) and after 3 months (T3). Besides NIHSS and mRS, stroke outcomes included the Barthel Index for the assessment of daily life activities and the Montreal Cognitive Assessment (MoCA). In case patients could not attend the T3 appointment at the University Hospital Essen due to rehabilitation or reduced health condition, telephone-based interviews were conducted to determine mRS. Death certificates were collected to confirm a mRS score of 6. All participants provided informed consent and protocols were approved by the institutional review board (2110271-BO).

### Legal issues for animal experimentation, randomization, and blinding

All experiments were performed with local government approval (Landesamt für Verbraucherschutz und Ernährung, Recklinghausen) in accordance to EU guidelines for the care and use of laboratory animals (Directive 2010/63/EU) and are reported in accordance to Animal Research: Reporting of In Vivo Experiments (ARRIVE) guidelines. Experiments were strictly randomized. The experimenters (D.Y., A.R.L.) performing the animal experiments, functional neurological studies, and histochemical studies were fully blinded at all stages of the study by other researchers (S.T., C.W.) preparing injectable solutions. These solutions with dummy names (A and B) were unblinded after termination of the study.

### Animal housing and sample size calculation

Mice were housed under a regularly reversed 12-hour light/12-hour dark cycle, allowing them unrestricted access to food and water. Animals were grouped in cages with 5 animals per cage. Animal surgeries and neurological tests were always performed in the morning hours throughout the study. Sample size planning was conducted with G*Power version 3.1.9.7 software. Experiments were scheduled for expected effect sizes that α errors of 0.05 and β errors of 0.2 were achieved.

### Intraluminal MCAO model

Focal cerebral ischemia was induced as described before ^15,45^. Male 10-12 week-old C57BL6/j mice (Envigo, Darmstadt, Germany) were anesthetized with 1.5% isoflurane (30% O_2_, remainder N_2_O), while 150 µl of buprenorphine (0.1 mg/kg; Reckitt Benckiser, Slough, U.K.) was injected subcutaneously for analgesia. Rectal temperature was measured and maintained at ∼37°C using a feedback-controlled heating system (Fluovac; Harvard apparatus, Holliston, MA, U.S.A.). In all studies, a flexible laser Doppler flow (LDF) probe was attached on the animals’ skulls above the core of the middle cerebral artery territory for regional cerebral blood flow recording (Perimed, Stockholm, Sweden). After a midline neck incision, the left common carotid artery (CCA), internal carotid artery (ICA), and external carotid artery (ECA) were exposed. The CCA and ECA were ligated by sutures, the ICA was temporarily clipped. A silicon-coated 7.0 nylon monofilament (Doccol, Sharon, MA, U.S.A.) was introduced into the CCA through a small incision and advanced up to the bifurcation of the middle cerebral artery. After 30 min of MCAO, the filament was withdrawn to initiate reperfusion. LDF recordings were continued over 15 min. In all studies, LDF recordings during MCAO and after reperfusion did not differ between groups. The LDF probe was finally removed and all wounds were carefully sutured. During the first 3 days post stroke, animals received daily i.p. injections of carprofen (4 mg/kg; Bayer Vital, Leverkusen, Germany) as anti-inflammatory drug.

### S. pneumoniae pneumonia

For bacterial lung infection, a pneumococcal serotype 1 strain (*Streptococcus pneumoniae* SV1, ATCC 33400) was used. Bacteria were grown overnight on Columbia blood agar plates (PB5039A; Thermo Scientific, Waltham, MA, U.S.A.) at 37°C. Single colonies were resuspended, cultured in 10 mL brain-heart infusion broth (TV5090E; Thermo Scientific) to mid-logarithmic phase (OD600=0.045-0.055; NanoDrop 1000 spectrophotometer; Thermo Scientific), and 800 µl of culture supplemented with 200 µl of 86 % glycerol was frozen at -80 °C. For infection, bacteria were again grown to mid-logarithmic phase, centrifuged at 1500g at 4°C for 10 min, and resuspended in 550 µl 0.01 M PBS. 50 µl suspension per animal corresponding to 1x10^8^ CFUs *S. pneumoniae* were used for infection ^46^.

Mice were anesthetized by intraperitoneal injection of ketamine/ xylazine (80 mg/kg and 10 mg/kg b.w., respectively). For intratracheal instillation, the mouse was placed on a 60° angled self-made stand and secured by a Velcro stip. The teeth of the mouse were suspended on a thin rubber band and the tongue was pulled out. A 22 gauge cannula with a blunted end was inserted into the mouse trachea with the help of a small-animal laryngoscope (73-4867; Harvard Apparatus, Holliston, MA, U.S.A.). The needle inside was removed and 50 µl of *S. pneumoniae* suspension was pipetted into the cannula. A MiniVent 845 ventilator (73-4867; Harvard Apparatus) was connected to the cannula to evenly spread the suspension into the lungs ^46^. In line with earlier studies ^46^, reproducible neutrophil and macrophage infiltrates were noted in the animals’ lungs on occasion of the animal sacrifice 48 hours post-pneumonia in MCAO pneumonia mice.

### Drug administration

As antibiotic, amoxicillin (15 mg/kg; Duphamox LA; Zoetis, Parsipanny, NJ, U.S.A.) dissolved in vehicle (Lipovenös MCT 20%; Fresenius Kabi, Bad Homburg, Germany) or an equivalent volume of vehicle was administered s.c. 3 hours post-pneumonia. This injection was repeated at 8 hour-intervals until 3 days post pneumonia (dpp), as previously reported ^47^. Thus, a total of 9 injections were applied.

To explore the role of neutrophils in the pathological sequelae associated with stroke-associated pneumonia, neutrophils were depleted by i.p. injection of 200 µg of anti-Ly6G antibody (clone 1A8; BioXCell, West Lebanon, NH, U.S.A.) 24 hours before pneumonia and 24 hours after pneumonia. This dose almost completely clears all neutrophils from the blood, as we previously showed ^15^. An equivalent amount of isotype control IgG (clone 2A3; BioXCell) was administered as control treatment.

To degrade circulating NET DNA, deoxyribonuclease-I (DNase-I; 11284932001; Roche, Basle, Switzerland) was applied at a dose of 10 µg i.v. and 50 µg i.p. 30 min before pneumonia. The i.p. dose (50 µg DNase-I) was repeated every 12 hours until animal sacrifice, according to previously published protocols ^48^. On each occasion, an equivalent amount of vehicle was injected as control treatment.

For NET formation blockade, the gasdermin-D inhibitor LDC7559 (HY-111674; MedChemExpress, Monmouth Junction, NJ, U.S.A.) dissolved in corn oil was injected i.p. at a dose of 10 mg/kg b.w. 30 min before pneumonia in animals sacrificed at 2 dpp, as previously reported ^19^. In animals sacrificed at 56 dpp, LDC7559 (10 mg/kg b.w.) was administered i.p. 30 min before pneumonia and at 1, 2, 4, and 7 dpp (animal set A) or immediately after MCAO, 24 hours post-MCAO, 30 min before pneumonia and at 1, 2, 4, and 7 dpp (animal set B). For the control group, an equivalent volume of vehicle (corn oil) was applied on each occasion.

To inhibit MPO, the pharmacological inhibitor AZD4831 (HY-145581; MedChemExpress) was administered by oral gavage at a dose of 10 µmol/kg, starting 2 hours before pneumonia induction and subsequently once every 24 hours until animal sacrifice, as previously published ^24^. AZD4831 was dissolved in normal saline containing 1% DMSO, 44% PEG-300, and 5% Tween-80. Control mice received an equal amount of vehicle solution.

To neutralize histone proteins, mice were treated i.p. with 20 µg anti-histone H2A/H4 antibody (clone BWA3; Creative Biolabs, Shirley, NY, USA) ^25^. For animals sacrificed at 2 dpp, injections were administered 30 min before pneumonia induction and 1 day later. For animals sacrificed at 56 dpp, injections were performed 30 minutes before pneumonia induction and at 1, 2, and 3 dpp. Control mice received 20 µg of isotype control IgG (clone MOPC-21; BioXCell, West Lebanon, NH, USA) i.p. at the same time points.

### Neurological score

The modified Clark score is a comprehensive score that examines both general and focal neurological deficits, for which a composite score is formed ^45^. This score was studied daily until 7 dpp, subsequently evaluated at 9 dpp, 11 dpp, and 14 dpp, followed by weekly assessments until 56 dpp.

### Tight rope test

The tight rope test consists of a 60-cm-long rope connected to two platforms. Animals were placed on the middle of the rope and the time until reaching one of the platforms or dropping off the rope was recorded ^45^. The maximum testing time was 60 s. Recorded times were translated into scores. Animals were trained three times per day for three days before MCAO. After a baseline evaluation before MCAO, animals were tested before pneumonia induction at 0 dpp, and then weekly until 56 dpp.

### Rotarod test

The rotarod is a motor-coordination test that is assessed on a rotating drum ^45^. Mice were positioned on the drum, which then began to accelerate from 4 rpm to 40 rpm. The time when the animal dropped off the drum was recorded with a maximum of 300 s for each testing session. The animals were trained 3 times a day for 3 days before the experiments. After a baseline assessment, the mice were evaluated from 0 dpp prior to the induction of pneumonia, followed by weekly assessments up to 56 dpp.

### Open field test

The open field test consists of a square arena (40x40x30 cm) subdivided into a center, four borders, and four corners, in which animals were placed near the wall and observed for 10 min for evaluating spontaneous motor activity ^45^. The total distance covered was tracked using ANY-maze software (Stoelting, Dublin, Ireland). The open field test was performed at 28 dpp and 56 dpp.

### Novel object recognition test

The novel object recognition test uses the same arena as the open field test for assessing recognition memory for presented objects. In a training phase, two identical objects were placed 10 cm away from the walls in opposite corners of the arena. Mice were allowed to explore these objects for 10 min. One hour later, one familiar object was replaced by a novel one of different shape and appearance. Mice were again allowed to explore the chamber for 5 min, while they were tracked using ANY-maze software. Exploration behavior was defined as sniffing, licking, or touching the object with the forepaws. The novel object discrimination index = (novel object exploration time - familiar object exploration time) / (novel object exploration time + familiar object exploration time) was determined.

### Quantification of citrullinated histone H3 associated plasma DNA

NETs were quantified in plasma samples obtained 48 hours post-pneumonia by retroorbital blood puncture using a previously described capture ELISA via the measurement of citrullinated histone-H3 (CitH3) associated with DNA ^19^. Anti-CitH3 antibody (5 µg/mL; ab5103, Abcam) was coated overnight in 50 µl volume at 4°C onto 96-well plates followed by 5% bovine serum albumin (BSA) blocking for 2 h on a shaker at 300 rpm. Coated wells were washed three times with 300 µl washing buffer followed by addition of 50 µl plasma and 80 µl incubation buffer (including peroxidase-labeled anti-DNA antibody) for 2 hours at 300 rpm (Cell Death ELISAPLUS, Cat. 11774425001, Roche). Then, the wells were washed three times with 300 µl incubation buffer. Following the addition of 100 µl peroxidase substrate to the wells, samples were incubated in the dark for 30 min at room temperature on a shaker at 300 rpm. Afterward, 100 µl ABTS peroxidase stop solution was added to the wells. Absorbance was measured at 405 nm and absorbance at 490 nm was subtracted. Absorbance values were presented as a relative increase to control.

### MPO activity measurement

MPO was determined using a colorimetric activity assay (EEA016; Thermo Scientific) in plasma samples obtained via retro-orbital puncture at 24 hours post-pneumonia according to the manufacturer’s instructions.

### Animal sacrifice and measurement of infarct volume, brain edema, atrophy, and hemorrhages

At indicated time-points, mice were transcardially perfused with ice-cold 0.01 M PBS followed by 4% paraformaldehyde in 0.01 M PBS during 3% isoflurane anesthesia (30% O_2_, remainder N_2_O). After overnight post-fixation in 4% paraformaldehyde in 0.01 M PBS at 4°C, brains were dehydrated using 15% and 30% sucrose, frozen, and stored at -80°C. Then, 20 µm thick coronal sections were collected at 1 mm intervals across the forebrain for cresyl violet staining. Sections were scanned and analyzed by ImageJ software (National Institute of Health, Bethesda, MD, U.S.A.). In animals sacrificed at 2 dpp, infarct volume was determined by subtracting the volume of healthy brain tissue of the ischemic hemisphere from that of the contralateral hemisphere ^15^. Brain edema was assessed as increase in volume of the ipsilateral brain hemisphere compared to that of the contralateral hemisphere divided by the volume of the contralateral hemisphere ^15^. In animals sacrificed at 56 dpp, brain tissue atrophy was evaluated by the volume reduction of the ipsilateral hemisphere relative to the contralateral hemisphere divided by the volume of the contralateral hemisphere ^45^.

To analyze brain hemorrhages, 20-µm-thick coronal sections harvested at various levels of the infarct (+1.0, ±0.0, -1.0, and -2.0 mm rostral to bregma) were incubated in 0.05% diaminobenzidine (DAB) solution (D5905; Sigma-Aldrich). This substrate, oxidized by erythrocyte peroxidases, yields a dark brown staining. The area and incidence of brain hemorrhages were assessed at all levels ^21^. The area values determined were extrapolated to volume values by multiplying areas with section thickness values and integrating the values measured across the brain, considering that one of fifty consecutive brain sections was evaluated.

### Immunohistochemistry

20-µm-thick coronal sections from the bregma level were rinsed with 0.01 M PBS and permeabilized with 0.3% Triton X-100 in 0.01 M PBS. Subsequently, the sections were incubated in 0.01 M PBS containing 10% normal donkey serum (D9663; Sigma-Aldrich) for 1 hour at room temperature. Following that, sections were incubated overnight at 4 °C with primary antibodies. The primary antibodies used were Alexa Fluor-594-conjugated polyclonal donkey anti-IgG (A-21203; Thermo Scientific), polyclonal goat anti-intercellular adhesion molecule (ICAM-1) (AF796; R&D Systems, Minneapolis, MN, U.S.A.), monoclonal rat anti-CD45 (05-1416; Merck Millipore, Darmstadt, Germany), polyclonal rabbit anti-collagen-IV (AB756P; Merck-Millipore), monoclonal rat anti-glycoprotein-Ibα (GP-Ibα) (M043-0; Emfret Analytics, Eibelstadt, Germany), and polyclonal rabbit anti-ionized calcium binding adaptor protein (Iba)-1 (019–19741; Wako Chemicals, Neuss, Germany) antibodies. Then, the samples were rinsed and labeled with appropriate secondary Alexa Fluor-594 or Alexa Fluor-488 antibodies for 1 hour at room temperature. Nuclei were counterstained with Hoechst 33342 (Thermo Scientific). Sections were coverslipped and scanned using an inverted microscope equipped with apotome optical sectioning (Axio Observer.Z1; Carl Zeiss, Oberkochen, Germany). Extravasated IgG and ICAM-1 were evaluated by integrated signal intensities. For CD45^+^ leukocytes and Iba-1^+^ microglia/ macrophages, three 300 μm × 300 μm ROIs were chosen in the ischemic striatum to analyze cell density. Microvascular thrombosis was assessed by calculating the density of collagen-IV^+^ microvessels containing GP-Ibα^+^ clots.

### Microglia/macrophage morphology analysis

A Leica SP8 confocal microscope (objective HC PL APO CS2 63×/1.30, Leica Microsystems, Wetzlar, Germany) was used to assess the three-dimensional morphology of Iba-1^+^ microglia/ macrophages in the three representative ROIs. Z-stacks with dimensions of 184.7 × 184.7 × 15 µm were acquired at 0.5 µm intervals. To analyze the 3D cell morphology, our in-house script 3DMorph, based on MATLAB (The MathWorks Inc., Natick, MA, U.S.A.), was utilized. In summary, automated threshold setting and segmentation were employed for the detection of cell objects. Following skeletonization, microglial ramification index and cell territory were obtained.

### Flow cytometry of leukocytes

At 2 dpp, blood samples were collected from the animals’ hearts, and ischemic brain hemispheres were collected from the same animals. For blood samples, single-cell suspensions were obtained after erythrocyte lysis and two washing steps. Brain tissue samples were smashed on a 70 μm cell strainer in culture dish, centrifuged with 37% Percoll solution, and after washing single-cell suspensions were obtained. These single-cell suspensions were stained for 30 min at 4 °C with antibody cocktails described in Extended Data Table 1. Subsequently, cell suspensions were analyzed using a Cytoflex flow cytometer (Beckman–Coulter, Brea, CA, U.S.A.) and FlowJo software V10 (Ashland, OR, U.S.A.). The gating strategy is outlined in Extended Data Fig. 1.

### Isolation of blood neutrophils for proteome analysis

Blood samples were subjected to erythrocyte lysis using a lysis solution (NH_4_Cl 150 mM, KHCO_3_ 10 mM, EDTA 0.1 mM, pH 7.3, 20 °C) with 1 mL used for each sample. The reaction was stopped with 11 mL of ice-cold (4°C) DMEM after 2 min. Then, the samples were centrifuged at 460 g at 4°C for 5 min and the supernatant was removed. Samples were washed with ice-cold (4°C) 0.1 M PBS thereafter. Following resuspension of the pellet and incubation with Fc-blocking antibodies (CD16/CD32; clone 2.4G2; BD Biosciences, New Jersey, U.S.A.) at 4°C for 15 min, samples were incubated with anti-Ly6G antibody (clone 1A8, BD Biosciences), anti-CD11b antibody (clone M1/70, eBioscience, Affymetrix, California, U.S.A.), and DAPI (BioLegend, San Diego, CA, U.S.A.). After washing and resuspension, CD11b^+^ Ly6G^+^ live neutrophils were sorted using a FACS Aria cell sorter (BD Biosciences, BD, New Jersey, U.S.), and the purity of cells was assessed (≥95%). The cells were then stored in protein lysis buffer and frozen in -80 °C until future use.

### Sample preparation for proteomics

The sorted neutrophils (typically 40,000–50,000) were extensively washed with PBS and resuspended in lysis buffer (50 mM Tris-HCl (pH 7.8), 150 mM NaCl, and 1% SDS) supplemented with complete mini-EDTA-free protease inhibitor (Roche, Penzberg, Germany). Each sample was incubated with 2.5 U benzonase nuclease (Merck-Millipore) for 7 min at room temperature. The proteins were reduced for 30 min at 37°C in 10 mM dithiothreitol (DTT) and alkylated in 30 mM iodoacetamide for 30 min at room temperature in the dark. After that, the proteins were precipitated with nine volumes of ethanol for 2 h at -80°C and centrifuged for 45 min at 20,000 g. The supernatant was removed, the pellet was dried and dissolved first in 1 µL of 6 M GuHCl and then in 29 µL of 50 mM ammonium bicarbonate buffer, pH 7.8, containing 2 mM CaCl_2_ and 50 ng of trypsin (sequencing grade, Promega), before incubation for 15 h at 37°C. The concentration of peptides was measured on a Nanodrop spectrophotometer at 205 nm. The enzymatic digestion was stopped by acidifying the sample to pH<2.5 with TFA.

### LC-MS/MS data acquisition

Chromatographic separation of peptides was achieved with a two-buffer system (buffer A: 0.1% FA in H_2_O, buffer B: 0.1% FA in 80% ACN) on a UHPLC system (VanquishTM neo, Thermo Scientific). Attached to the UHPLC was a peptide trap (100 µm x 20 mm, 100 Å pore size, 5 µm particle size, C18, Nano Viper, Thermo Scientific) for online desalting and purification, followed by a 25 cm C18 reversed-phase column (75 µm x 250 mm, 130 Å pore size, 1.7 µm particle size, peptide BEH C18, nanoEase, Waters). Peptides were separated using a 70 min protocol with linearly increasing ACN concentration from 2.5 % to 37.5 % ACN over 60 min.

MS/MS measurements were performed on a quadrupole-orbitrap hybrid mass spectrometer (Exploris 480, Thermo Scientific). Eluting peptides were ionized using a nano-electrospray ionization source (nano-ESI) with a spray voltage of 1,800 and analyzed in data dependent acquisition (DDA) mode. For each MS1 scan, ions were accumulated for a maximum of 25 milliseconds or until a charge density of 3 x 10^6^ ions (AGC Target) was reached. Fourier-transformation based mass analysis of the data from the orbitrap mass analyzer was performed covering a mass range of m/z 350 – 1,400 with a resolution of 60,000 at m/z = 200. Peptides being responsible for the 20 highest signal intensities per precursor scan with a minimum intensity of 8 x 10^3^ and charge state from +2 to +6 were isolated within a m/z 2 isolation window and fragmented with a normalized collision energy of 30% using higher energy collisional dissociation (HCD). MS2 scanning was performed, covering a mass range starting at m/z 120 and accumulated for 50 ms or to an AGC target of 5 x 10^4^ at a resolution of 15,000 at m/z = 200. Already fragmented peptides were excluded for 30 s.

### Raw data processing

LC-MS/MS were searched with the Sequest algorithm integrated in the Proteome Discoverer software (v 3.0.0.757, Thermo Scientific) against a reviewed Mus musculus database obtained in December 2022. Carbamidomethylation was set as fixed modification for cysteine residues and the oxidation of methionine as well as methionine-loss and acetylation of the protein N-terminus were allowed as variable modifications. A maximum number of 2 missing tryptic cleavages was set. Peptides between 6 and 144 amino acids where considered. A strict cutoff (FDR<0.01) was set for peptide and protein identification. Quantification was performed using the Minora Algorithm, implemented in Proteome Discoverer. Results were filtered for high confident peptides, with enhanced peptide and protein annotations. Protein abundances were exported for subsequent statistical analysis and normalized.

### Data analysis, statistics, and visualization

Statistical analyses were conducted using GraphPad Prism version 9.5.1 for Windows (GraphPad Software, San Diego, California, U.S.A.). LDF recordings, neurological deficits, and behavioral tests were analyzed by repeated measures ANOVA, with subsequent LSD posthoc test. These longitudinal data were presented as mean ± standard deviation (SD). For histochemical and proteomics data that were normally distributed, two-tailed independent samples t-tests or one-way analysis of variance (ANOVA) followed by LSD posthoc tests were utilized, as appropriate. For data that did not follow a normal distribution, the Kruskal-Wallis test followed by Dunn’s tests was used. These cross-sectional data were presented in box plots containing median and mean values ± interquartile ranges (IQR) with minimum and maximum data as whiskers. Hemorrhage incidence data were assessed by Fisher’s exact test and reported as percent rates. P values < 0.05 were considered to indicate statistical significance.

LC-MS/MS data were filtered by removing samples with low abundance and proteins that were not found in at least 3 samples of one group and subsequently normalized using loess normalization (R-limma) followed by imputation with a uniform random 5th-10th percentile approach for missing values. Statistical significance was tested using Student t-test (scipy). Effect sizes were computed with signal-to-noise ratio (SNR) and log_2_ fold change (log_2_FC) where the log_2_FC is determined as 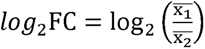, with 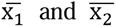 as the mean values, and the SNR is calculated using 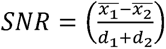 with d1 and d2 as the respective standard deviation. Significant regulation was considered for the proteins with an absolute log_2_FC >0.263 and p value <0.05. Heatmaps, volcano plots, and principal component analysis (PCA) were generated using Morpheus (https://software.broadinstitute.org/morpheus), GraphPad Prism version 9.5.1 for Windows, the Python package matplotlib (v.3.8.1) and seaborn (v.0.13.2).

## Acknowledgments

We thank the Institute of Experimental Immunology and Imaging center of University Hospital Essen for assistance in imaging studies.

## Author contributions

DMH and MG designed the experiments. DY and ARL, together with AMY, ST, and TT, performed the experiments. EP and JJ conducted neutrophil isolation. OS, BS, HS and DS performed proteomic data acquisition and analysis. DRE supervised the proteomic dataset. JG, YZ, CG, HT, MF, YL, and FJ collected clinical data. CW and NH provided advice on the mouse model. BK, BF, AAT, and VS provided technical support. DMH, DY, and ARL wrote the original draft of the manuscript. BS, HS, ED, JM, LK, BG, and OSh reviewed and edited the original draft. All authors contributed to editing and approved the final version of the manuscript.

## Data Availability Statement

The mass spectrometry data have been deposited to the ProteomeXchange Consortium via the PRIDE partner repository with the dataset identifier PXD053004 and will be publicly made accessible upon publication.

