## Supplementary material for "Histone neutralization protects the ischemic brain against stroke-associated pneumonia": Supplementary data_28.02.2026.docx

**Legends to Extended Data figures:**

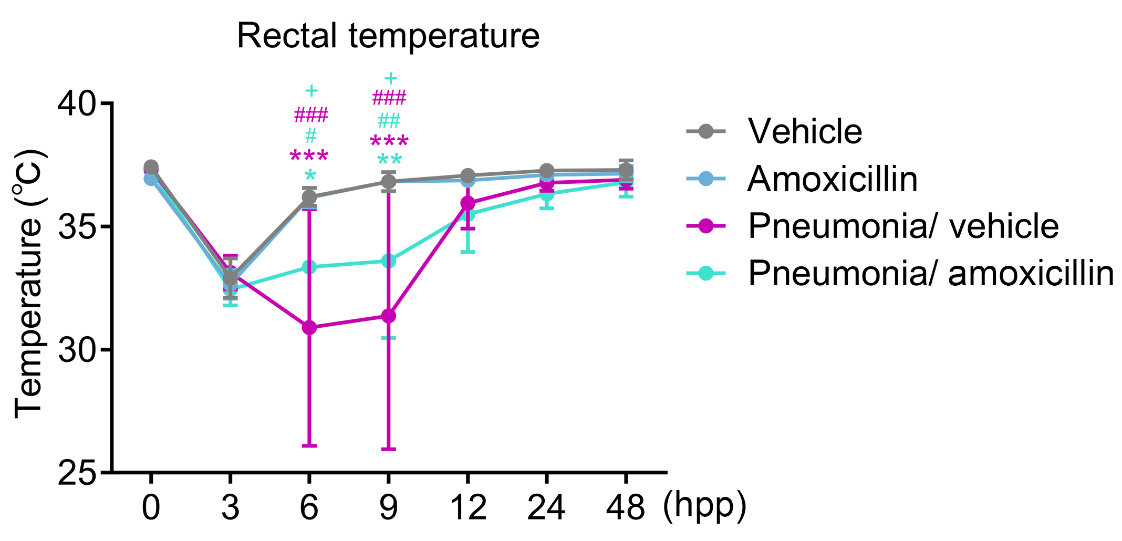

**Extended Data Fig.1. *S. pneumoniae* pneumonia induces post-ischemic hypothermia, which is attenuated by amoxicillin.** Rectal temperature of mice exposed to transient intraluminal middle cerebral artery occlusion (MCAO), followed by no pneumonia or pneumonia induced by intratracheal *S. pneumoniae* instillation (1x10^8^ CFUs) at 3 days post-MCAO and vehicle or amoxicillin (15 mg/kg t.i.d., s.c.) treatment starting 3 hours thereafter. *p<0.05, **p<0.01, ***p<0.001 compared with no pneumonia/ vehicle; ^#^p<0.05, ^##^p<0.01, ^###^p<0.001 compared with no pneumonia/ amoxicillin; ^+^p<0.05 compared with pneumonia/ vehicle (n=4 mice/ group).

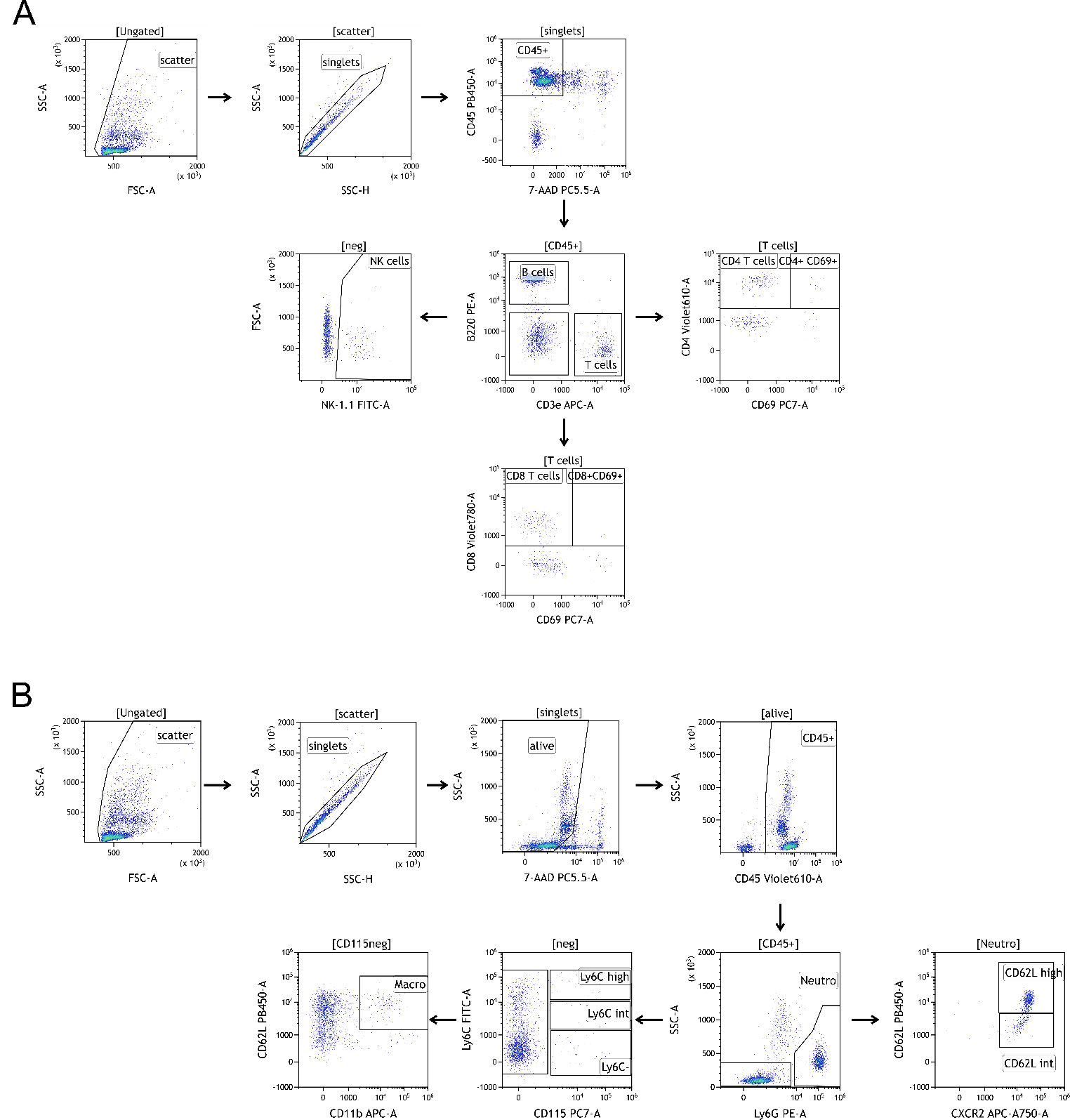

**Extended Data Fig.2. Gating strategy used for analyzing blood leukocytes and leukocyte activation by flow cytometry.** In two sets of studies, **(A)** lymphoid and **(B)** myeloid cells were examined in mice exposed to MCAO followed by *S. pneumoniae* pneumonia.

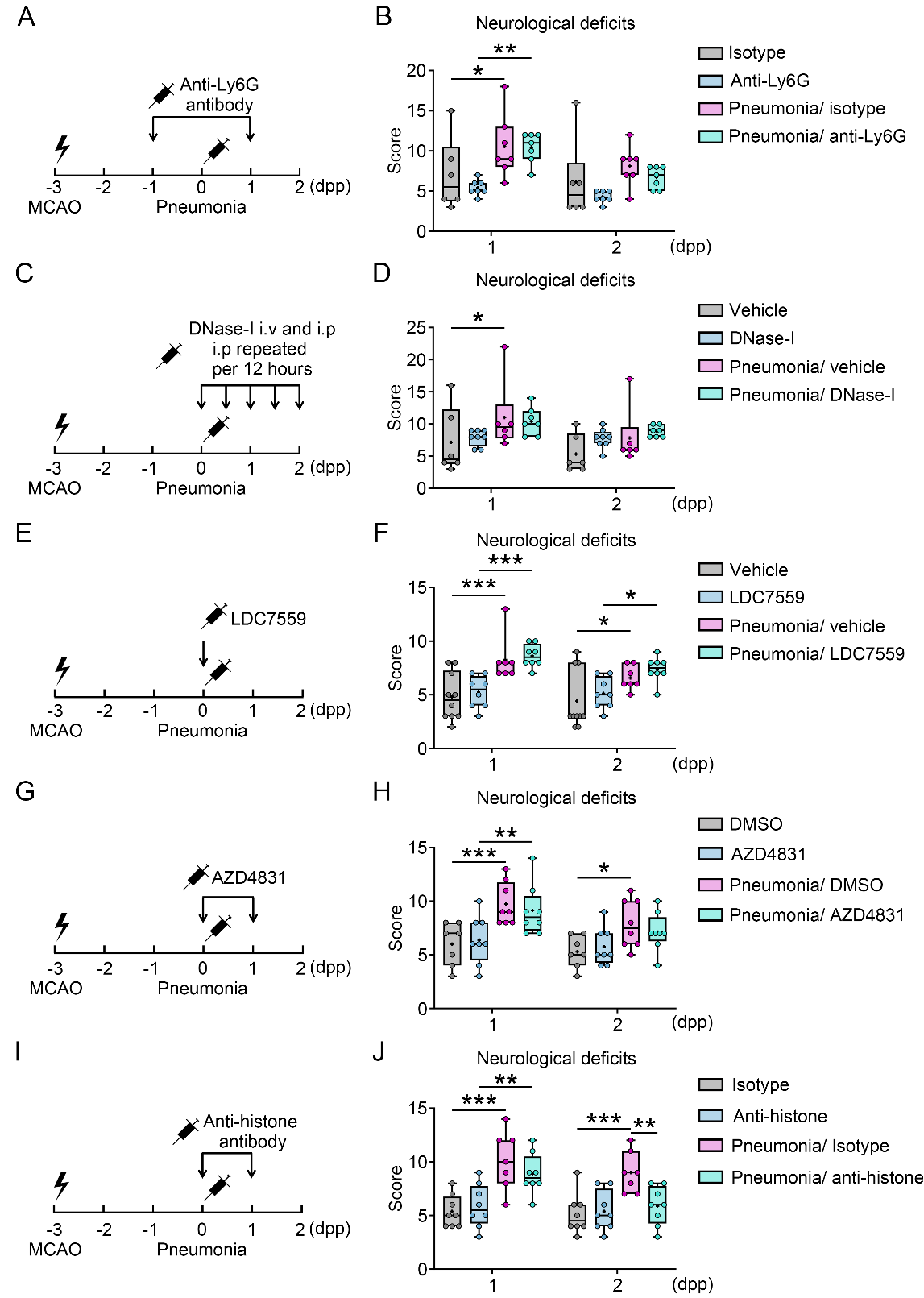

**Extended Data Fig.3. Effects of antibody-mediated neutrophil depletion, neutrophil extracellular DNA trap (NET) degradation, NET formation blockade, myeloperoxidase (MPO) inhibition, and histone neutralization on post-ischemic neurological deficits. (A, C, E, G, I)** Time-line of animal experiments and **(B, D, F, H, J)** neurological deficits evaluated by a comprehensive behavioral score of MCAO mice without pneumonia or MCAO mice exposed to *S. pneumoniae* pneumonia at 3 days post-MCAO, which were treated with **(A, B)** isotype antibody or anti-Ly6G antibody (1A8; 200 µg i.p.), which depletes neutrophils (*15*), 1 day before and 1 day after pneumonia (i.e., at 2 and 4 days post-MCAO), **(C, D)** vehicle or DNase-I (10 µg i.v. and 50 µg i.p. 30 min before pneumonia, followed by 50 µg b.i.d.), **(E, F)** vehicle or the gasdermin-D inhibitor LDC7559 (10 mg/kg i.p.), which prevents NET formation (*19*), 30 min before pneumonia and 1 day after pneumonia, **(G, H)** vehicle or the MPO inhibitor AZD4831 (10 µmol/kg via oral gavage) 2 hours before and 1 day after pneumonia, or **(I, J)** isotype IgG or neutralizing anti-histone antibody (BWA3; 20 µg i.p.) 30 min before and 1 day after pneumonia, followed by animal sacrifice at 2 dpp (that is, 5 days post-MCAO). *p<0.05, **p<0.01, ***p<0.001 (n=6-10 mice/ group).

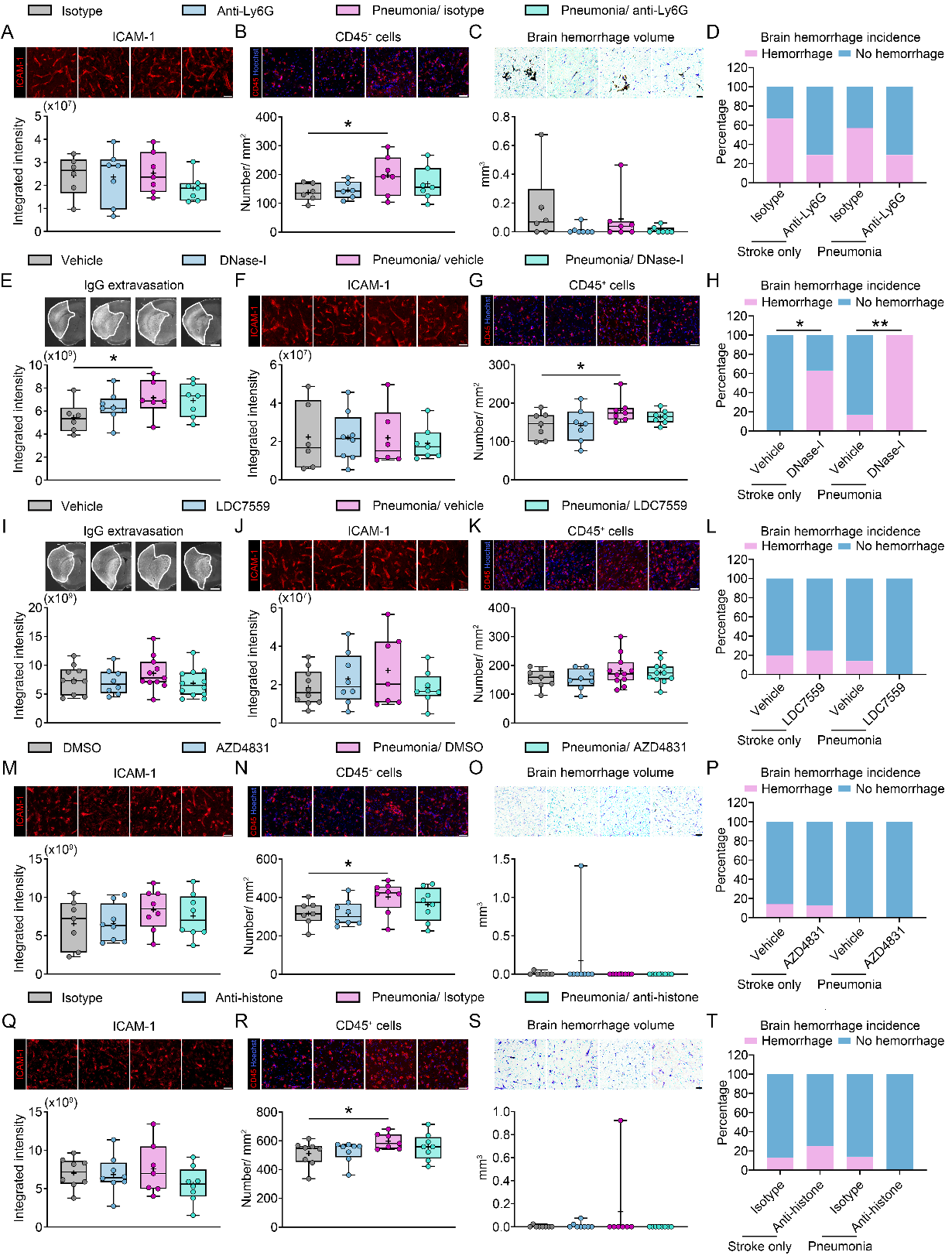

**Extended Data Fig.4. Effects of neutrophil depletion, NET degradation, NET formation blockade, MPO inhibition, and histone neutralization on cerebral microvascular ICAM1 abundance, brain leukocyte infiltrates, brain hemorrhage formation, and IgG extravasation. (A, F, J, M, Q)** Cerebral microvascular ICAM1 abundance, **(B, G, K, N, R)** brain-infiltrating CD45+ leukocyte density, **(C, O, S)** brain hemorrhage volume, **(D, H, L, P, T)** hemorrhage incidence, and **(E, I)** IgG extravasation in the previously ischemic brain tissue of MCAO mice without pneumonia or MCAO mice exposed to *S. pneumoniae* pneumonia at 3 days post-MCAO, which were treated with **(A-D)** isotype antibody or anti-Ly6G antibody (1A8; 200 µg i.p.), which depletes neutrophils (*15*), 1 day before and 1 day after pneumonia (i.e., at 2 and 4 days post-MCAO), **(E-H)** vehicle or DNase-I (10 µg i.v. and 50 µg i.p. 30 min before pneumonia, followed by 50 µg b.i.d.), **(I-L)** vehicle or the gasdermin-D inhibitor LDC7559 (10 mg/kg i.p.), which prevents NET formation (*19*), 30 min before pneumonia and 1 day after pneumonia, **(M-P)** vehicle or the MPO inhibitor AZD4831 (10 µmol/kg via oral gavage) 2 hours before and 1 day after pneumonia, or **(Q-T)** isotype IgG or neutralizing anti-histone antibody (BWA3; 20 µg i.p.) 30 min before and 1 day after pneumonia, followed by animal sacrifice at 2 dpp (that is, 5 days post-MCAO). Representative immunohistochemistry images are shown. *p<0.05, **p<0.01 (n=6-10 mice/ group). Scale bars: 50 µm (in **(A, B**, **F, G**, **J, K**, **M, N, Q, R)**; 50 µm (in **(C, O, S)**; 1 mm (in **(E, I)**).

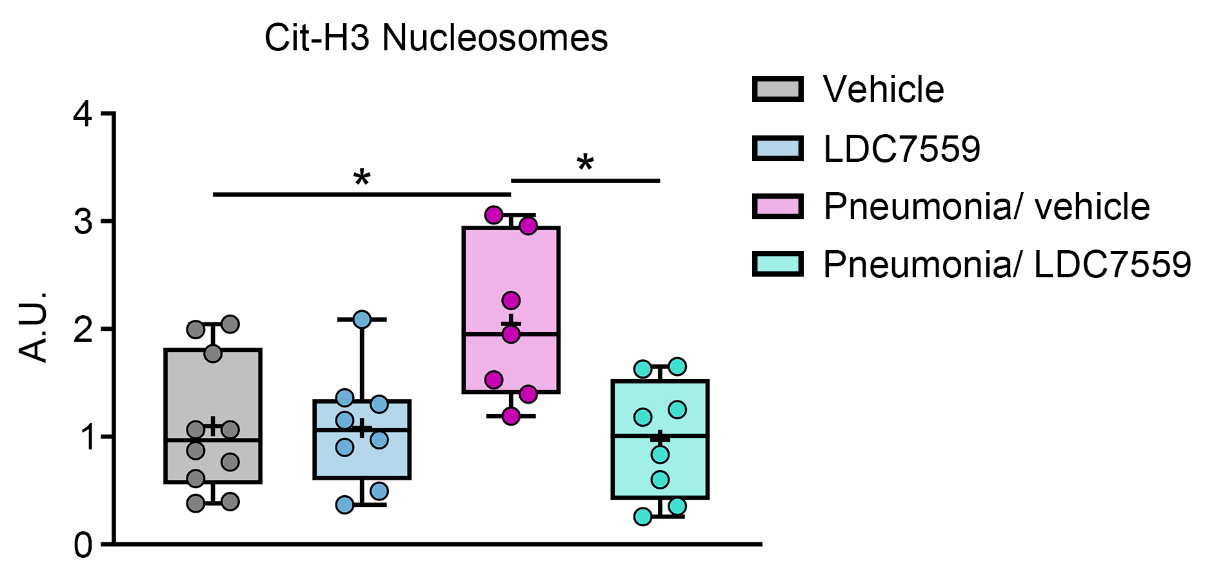

**Extended Data Fig.5. Effects of NET formation blockade by LDC7559 on DNA-associated citrullinated histone H3 levels in the blood.** Citrullinated histone-H3 (CitH3) protein associated with DNA assessed by capture ELISA in plasma samples obtained 48 hours post-pneumonia (120 hours post-MCAO) from MCAO mice without pneumonia or MCAO mice exposed to *S. pneumoniae* pneumonia at 3 days post-MCAO, which were treated with vehicle or the gasdermin-D inhibitor and NET formation blocker LDC7559 (10 mg/kg i.p.) 30 min before pneumonia. *p<0.05 (n=7-10 mice/ group).

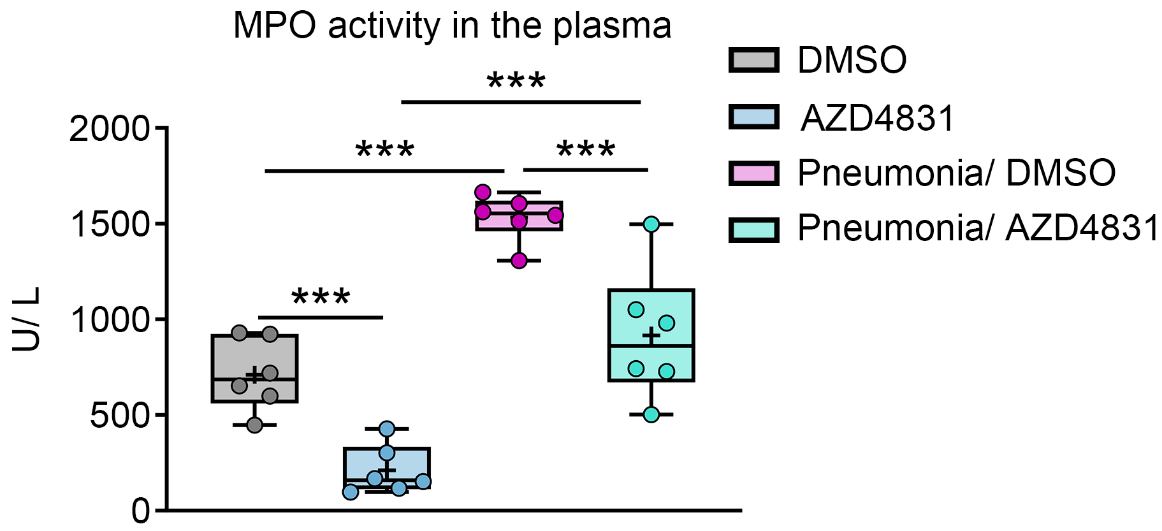

**Extended Data Fig.6. Effects of MPO inhibitor AZD4831 on MPO activity in the blood.** MPO activity determined using a colorimetric activity assay in plasma samples obtained 24 hours post-pneumonia (96 hours post-MCAO) from MCAO mice without pneumonia or MCAO mice exposed to *S. pneumoniae* pneumonia at 3 days post-MCAO, which were treated with vehicle or the MPO inhibitor AZD4831 (10 µmol/kg via oral gavage) 2 hours before pneumonia. ***p<0.001 (n=6 mice/ group).

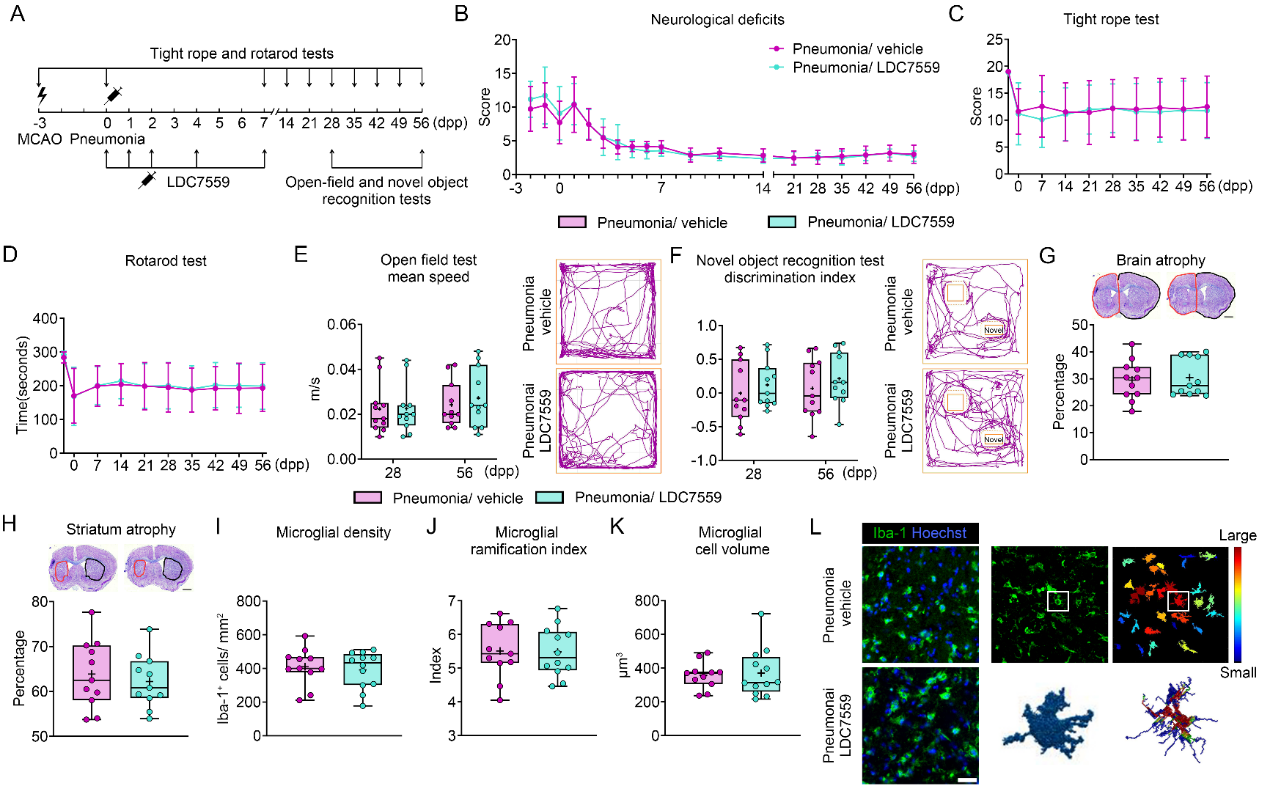

**Extended Data Fig.7. NET formation blockade does not influence long-term neurological outcome and brain atrophy, when initiated on occasion of pneumonia. (A)** Time-line of animal experiments. Motor-coordination deficits evaluated using **(B)** the neurological score, **(C)** tight rope tests, and **(D)** RotaRod tests, **(E)** spontaneous motor activity assessed by open field tests (tracking paths of representative animals are also shown), **(F)** novel object recognition assessed by object recognition tests (tracking paths of representative animals are also shown), **(G)** whole brain atrophy and **(H)** striatum atrophy, assessed by cresyl violet staining, and **(I-L)** density and activation of Iba1^+^ microglia in the ischemic striatum, assessed by cell counting and morphology analysis following microglia segmentation, skeletonization, and reconstruction, of MCAO mice not exposed to pneumonia or MCAO mice exposed to *S. pneumoniae* pneumonia at 3 days post-MCAO, which were treated with vehicle or the NET formation blocker LDC7559 (10 mg/kg i.p.) starting on occasion of pneumonia, followed by animal sacrifice at 56 dpp (behavioral and histochemical studies in **(B-H)**) or animal sacrifice at 2 dpp (microglial density and morphology studies in **(I-L)**). Representative images in **(G, H, L)** are depicted. No significant differences were noted between groups (n=11-12 mice/ group). Scale bars: 1 mm (in **(G, H)**); 30 μm (in **(L)**).

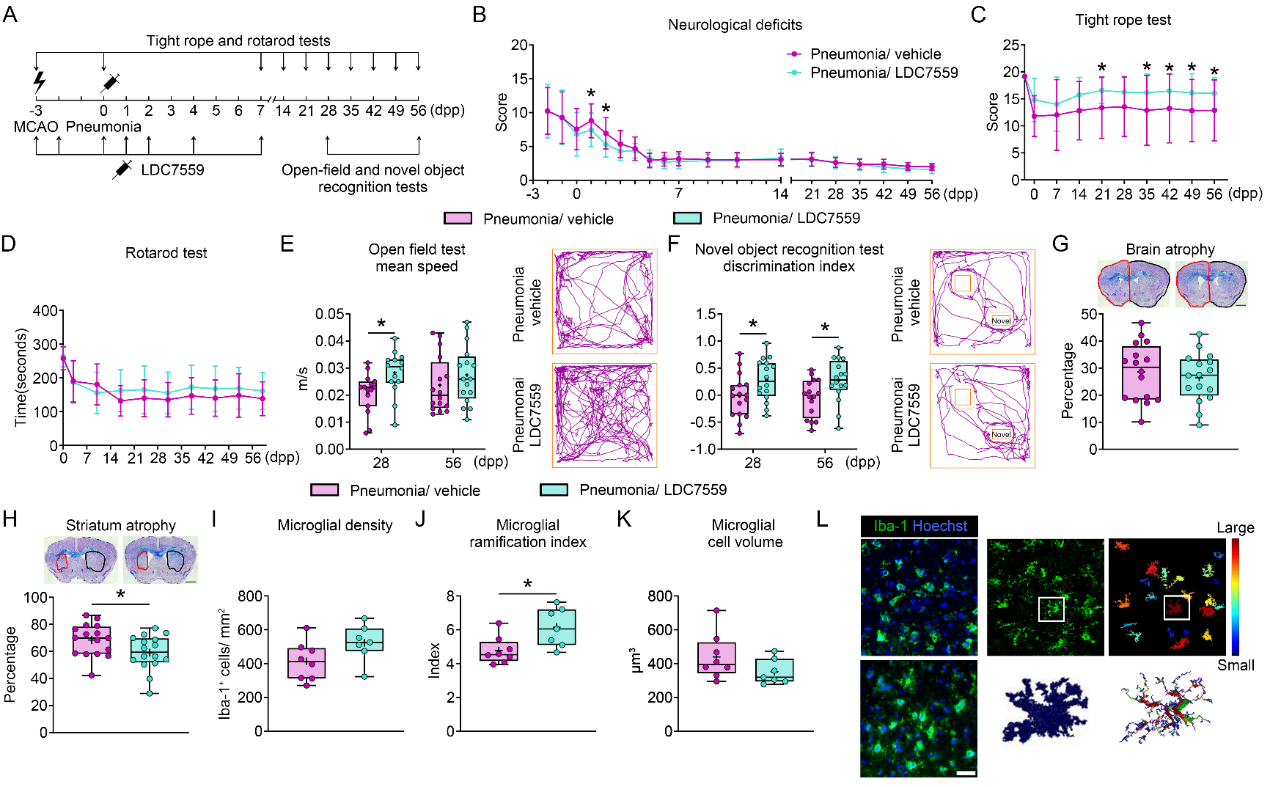

**Extended Data Fig.8. NET formation blockade by LDC7559 enhances long-term neurological recovery, reduces microglial activation, and reduces striatum atrophy, when initiated immediately after MCAO. (A)** Time-line of animal experiments. Motor-coordination deficits evaluated using **(B)** the neurological score, **(C)** tight rope tests, and **(D)** RotaRod tests, **(E)** spontaneous motor activity assessed by open field tests (tracking paths of representative animals are also shown), **(F)** novel object recognition assessed by object recognition tests (tracking paths of representative animals are also shown), **(G)** whole brain atrophy and **(H)** striatum atrophy, assessed by cresyl violet staining (samples of representative animals are also shown), and **(I-L)** density and activation of Iba1^+^ microglia in the ischemic striatum, assessed by cell counting and morphology analysis following microglia segmentation, skeletonization, and reconstruction, of MCAO mice not exposed to pneumonia or MCAO mice exposed to *S. pneumoniae* pneumonia at 3 days post-MCAO, which were treated with vehicle or the NET formation blocker LDC7559 (10 mg/kg i.p.) immediately after MCAO, followed by animal sacrifice at 56 dpp (behavioral and histochemical studies in **(B-H)**) or animal sacrifice at 2 dpp (microglial density and morphology studies in **(I-L)**). Representative images in **(G, H, L)** are depicted. *p<0.05 (n=16 mice/ group). Scale bars: 1 mm (in **(G, H)**); 30 μm (in **(L)**).

**Extended Data Table 1. Antibodies used for flow cytometry**

| Antigen | Conjugate | Host/isotype | Clone | Supplier |
| --- | --- | --- | --- | --- |
| Mouse CD45 | Pacific blue | Rat IgG2b, kappa | 30F11 | BioLegend |
| Mouse CD45 | BV 605 | Rat IgG2b, kappa | 30F11 | BioLegend |
| Mouse Ly6G | Phycoerythrin (PE) | Rat IgG2a, kappa | 1A8 | BioLegend |
| Mouse CXCR2  (CD182) | APC/Cyanine7 | Rat IgG2a, kappa | SA044G4 | BioLegend |
| Mouse CD62L | eFluor 450 | Rat IgG2a, kappa | MEL-14 | eBioscience |
| Mouse Ly6C | Fluorescein isothiocyanate (FITC) | Rat IgM, kappa | AL21 | BD Biosciences |
| Mouse CD11b | Allophycocyanin (APC) | Rat IgG2b, kappa | M1/70 | eBioscience |
| Mouse CD115 | PE-Cy7 | Rat IgG2a, kappa | AFS98 | eBioscience |
| Mouse CD3ε | Alexa Fluor 647 | Hamster IgG | 145-2C11 | BioLegend |
| Mouse CD4 | BV 605 | Rat IgG2a, kappa | RM4-5 | BD Biosciences |
| Mouse CD8 | BV 786 | Rat IgG2a, kappa | 53-6.7 | BD Biosciences |
| Mouse B220 | PE | Rat IgG2a, kappa | RA3-6B2 | BD Biosciences |
| Mouse NK-1.1 | FITC | Rat IgG2a, kappa | PK136 | BD Biosciences |
| Mouse CD69 | PE-Cy7 | Hamster IgG | H1.2F3 | BioLegend |
